# Rab7 deficiency induces lysosome formation from recycling endosomes leading to an increased degradation of cell surface proteins

**DOI:** 10.1101/2024.05.13.593900

**Authors:** Guan M. Wang, Peng Xu, Kaikai Yu, Shiny Shengzhen Guo, Reinhard Fässler

## Abstract

Cell surface receptors such as integrins are repeatedly internalized from and recycled back to the plasma membrane before routed to lysosomes for degradation. In search for modulators of β1 integrin surface stability, we identified the Rab7 small GTPase, believed to be required for lysosome biogenesis, as integrin stabilizer. We show that Rab7 deficiency produces late endosomes and lysosomes with acidic pH, lysosome-specific proteins and membrane architectures that are functional in protein degradation and organelle fusion. Furthermore, Rab7-deficient lysosomes form from Rab4- and transferrin receptor-positive recycling endosomes, resulting in the degradation of proteins designated for recycling. Finally, we also found that overexpression of Rab4 can direct lysosome formation from recycling endosomes in absence as well as presence of Rab7, however, the latter to a much lesser extent. Our findings reveal a lysosome biogenesis and lysosomal protein degradation pathway that becomes dominant in absence of Rab7 or when Rab4 is highly abundant.

## Introduction

Eukaryotic cells have a highly dynamic endomembrane system that compartmentalizes biochemical reactions and exchanges a multitude of molecules with the extracellular environment by means of endo- and exocytosis. The organization of the endomembrane system depends on an accurately regulated trafficking of membranes, which is orchestrated by the evolutionarily conserved superfamily of small Rab GTPases (Homma et al., 2021; Stenmark, 2012; Wandinger-Ness & Zerial, 2014) that cycle between an active, GTP-bound state and an inactive, GDP-bound state. The activity state of Rab proteins dictates their localization to specific endomembranes such as ER, Golgi, endosomes and lysosomes, and the recruitment of effector proteins that regulate membrane trafficking resulting in the biogenesis, transport, tethering and fusion of vesicles and organelles.

Integrins are ubiquitously expressed, mediate cell-extracellular matrix (ECM) and cell-cell adhesion and are α/β heterodimeric type I transmembrane receptors (Hynes, 2002) that, once synthesized and exocytosed to the cell surface, are recycled numerous times from and back to the plasma membrane by the endosomal system before being eventually degraded by lysosomes. Integrin trafficking is controlled by several Rab GTPases family members including but not limited to Rab5, Rab4, Rab11 and Rab7 (Moreno-Layseca et al., 2019; Paul et al., 2015). Rab5 governs the fusion of endocytic vesicles with the early endosomes as well as homotypic fusion of early endosome, leading to the delivery of internalized integrins and numerous other proteins to the intracellular sorting center (Zeigerer et al., 2012; Zerial & McBride, 2001). Rab4, which is also present on early endosomes, mediates cargo recycling either directly from early endosomes back to the plasma membrane or via dedicated protein recycling organelles, including the recycling endosomes (Rab4^+^ and Rab11^+^) and the Rab11^+^ perinuclear recycling compartment (PNRC) (Moreno-Layseca et al., 2019; Sonnichsen et al., 2000). Transmembrane proteins destined for degradation are sorted from the endosomal limiting membrane into the endosomal lumen via the budding of intraluminal vesicles (ILVs) (Babst, 2011). Endosomes with gradually accumulating ILVs are known as multivesicular bodies (MVBs) or late endosomes. The transition from early endosome to late endosomes is accompanied by the exchange of Rab5 by Rab7, acidification of the lumen, and changes in lipid and protein composition of the endosomal membrane (Huotari & Helenius, 2011; Langemeyer et al., 2018; Saftig & Klumperman, 2009). Under the control of Rab7, late endosomes mature into endolysosomes by acquiring lysosomal proteins from transport carriers and finally into lysosomes, where transmembrane proteins including integrins are degraded (Guerra & Bucci, 2016; Langemeyer et al., 2018).

Whereas the three Rab5 isoforms (Rab5A, Rab5B, and Rab5C) or the two Rab4 isoforms (Rab4A and Rab4B) share similar subcellular localizations and functions (Bucci et al., 1995; Homma et al., 2021; Homma et al., 2019; Perrin et al., 2013), only Rab7A is referred to as Rab7, because Rab7B shares limited similarity and no functional overlap with the evolutionarily conserved Rab7A (Yang 2004, Progida 2010, Mackiewicz and Wyroba 2009). Rab7A (from now on called Rab7) is ubiquitously expressed, controls the membrane protein and lipid composition of the endo-lysosomal system, and is therefore considered as the master regulator of lysosome biogenesis (Guerra & Bucci, 2016; Langemeyer et al., 2018), whereas Rab7B is expressed in few tissues and controls endosome-to-Golgi transport (Progida et al., 2010; Yang et al., 2004). Although the *Rab7* gene deletion results in embryonic lethality in mice (Kawamura et al., 2012; Roy et al., 2013), frogs (Kreis et al., 2021), flies (Cherry et al., 2013), worms (Skorobogata & Rocheleau, 2012), as well as protozoan parasites (Silverman et al., 2011), Rab7 deficiency does not impair viability of mammalian cell lines cultured *in vitro*. The grave consequences of Rab7 loss *in vivo* opposed to the normal cell viability *in vitro* points to unapparent but severe cellular dysfunction(s). So far, lack of Rab7 has been associated in mammalian cells with impaired autophagy (Kuchitsu & Fukuda, 2018; Kuchitsu et al., 2018; Roy et al., 2013) and loss of the Rab7 ortholog Ypt7 in yeast with impaired vacuole (yeast lysosome) biogenesis and impaired degradation (Haas et al., 1995; Wichmann et al., 1992).

In the present study, we carried out an unbiased whole genome screen in the human haploid cell line HAP1 to identify novel genes that regulate the stability of integrin cell surface levels. We found that Rab7 loss massively decreased rather than increased total and cell surface levels of integrins as well as other cell membrane proteins. In search for a mechanistic explanation for this striking finding, we found a Rab7-independent pathway that becomes activated in the absence of Rab7 expression and generates lysosomes from Rab4- and transferrin receptor-containing recycling endosomes.

## Results

### *RAB7A* gene stabilizes Itgb1 levels

To identify novel regulators of β1 integrin (Itgb1, encoded by the *ITGB1* gene in human and the *Itgb1* gene in mouse) trafficking we performed an unbiased genome-wide insertional mutagenesis screen in the human haploid cell line HAP1 using gene-trapping retroviruses carrying a splice acceptor site followed by the cDNA encoding the green fluorescent protein (GFP) (Carette et al., 2009; Jae et al., 2013). We generated 2.5 x 10^9^ mutagenized HAP1 cells, which were fixed, stained with monoclonal anti-Itgb1 antibody and sorted by flow cytometry to obtain the 5% cells with the lowest and the 5% cells with the highest Itgb1 surface levels (Figure 1A). Next, we determined the sites of gene-trap insertions by next-generation sequencing (NGS) and counted the total number of gene mutations in the Itgb1-low and -high cell populations, which revealed a high number of disruptive mutations in both cell populations. In the Itgb1-low population, we found known regulators of Itgb1 surface levels, such as *SNX17* and the SNX17-associated retriever complex consisting of *VPS26C*, *VPS29,* and *VPS35L* (Figure 1B-C; indicated with orange dots). Expectedly, we also identified mutations in genes coding for binding partners of the retriever complex including the CCC, WASH and Arp2/3 complexes (Figure 1B-C; indicated with blue dots), and genes required for Itgb1 maturation including the α integrin subunits that heterodimerize with Itgb1 (*ITGA2*, *ITGA4*, *ITGA6,* and *ITGAE*), proteins of the ER translocation machinery (*SEC62*), enzymes responsible for Itgb1 glycosylation (*GANAB*, *SRD5A3*, and *OSTC*) and chaperones (*HSP90B1*) (Figure 1B-C; indicated with green dots). Against all expectations, however, we also identified *RAB7A* as a potent stabilizer of Itgb1 surface levels (Figure 1B-C; indicated with red dot), which is in stark contrast to Rab7’s established role as master regulator of lysosome biogenesis/maturation and protein degradation. We decided to investigate how loss of Rab7 expression decreases Itgb1 surface levels.

**Figure 1.**
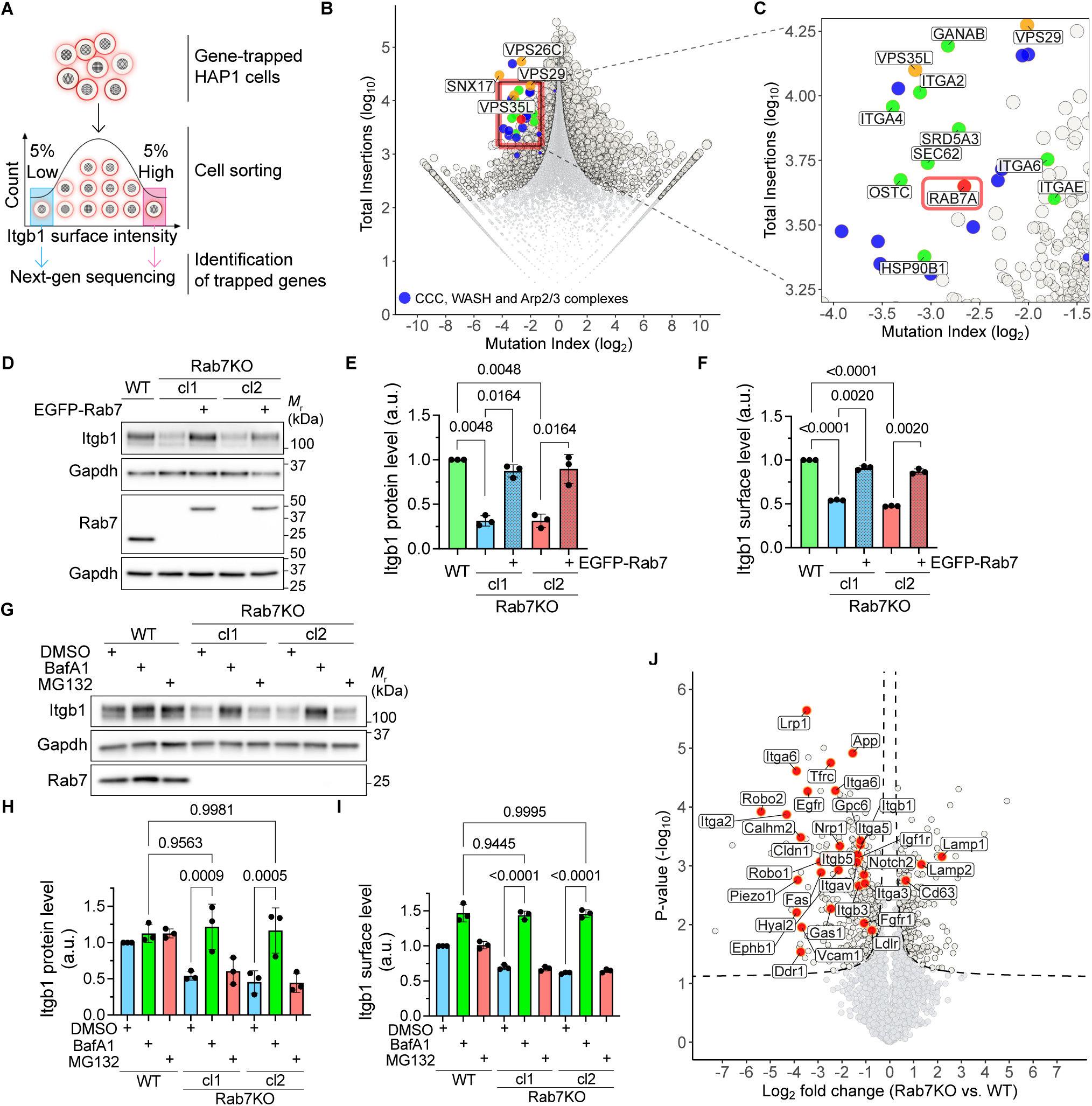
Rab7 deficiency decreases β1 integrin protein levels. **(A)** Schematic overview of the haploid genetic screen. The 5% lowest (Itgb1^LO^) and the 5% highest Itgb1 (Itgb1^HI^) surface levels were FACS-sorted and analyzed for gene trap insertion. **(B)** Haploid genetic screen for Itgb1^HI^ and Itgb1^LO^ surface levels. In the fishtail plot, genes enriched in the Itgb1 high and low populations are colored in blue and apricot, respectively. Dots represent individual genes and dot size corresponds to false discovery rate (FDR)-adjusted p-values (P_adj_) calculated with the Chi-square test. The y-axis indicates the number of disruptive insertions per gene and the x-axis indicates the mutation index (MI), which describes the frequency of independent insertions in Itgb1^HI^ channel over the frequency of insertions in the Itgb1^LO^ channel for each gene. Dark grey dots indicate genes with significant enrichment of insertions (P_adj_<10^-10^) and light grey dots with insignificant enrichment (P_adj_>10^-10^). Blue dots indicate genes coding for components of the retriever complex, orange dots indicate genes coding for components of the CCC, WASH and Arp2/3 complex, red dot indicate the Rab7 gene. **(C)** Close-up of highlighted region in **(B)**. **(D, E)** WB (D) and densitometric quantification (E) of Itgb1 in WT and Rab7KO fibroblasts, and Rab7KO fibroblasts stably re-expressing EGFP-Rab7. Gapdh served as loading control. Rab7KO cl.1 and cl.2 are independently generated cell clones. In WB (D) Itgb1 appears as 100 kDa immature and 125 kDa mature protein due to different glycosylation. The latter was quantified (E). Statistics was analyzed by two-sided multiple paired *t*-test with Holm-Šidák correction. Data are shown as Mean±SD, n=3 independent experiments. **(F)** Cell surface levels of Itgb1 on indicated cell lines determined by flow cytometry. Statistical tests were carried out as in E. Data are shown as Mean±SD, n=3 independent experiments. **(G, H)** WB (G) and densitometric quantification (H) of Itgb1 in WT and Rab7KO fibroblasts treated with DMSO (0.1% v/v), the lysosome inhibitor Bafilomycin A1 (BafA1, 10nM) or the proteasome inhibitor MG132 (100nM). Gapdh served as loading control. Statistics was analyzed by ordinary one-way ANOVA with Šidák’s post hoc tests. Data are shown as Mean±SD, n=3 independent experiments. **(I)** Cell surface levels of Itgb1 on indicted cell lines treated with DMSO, BafA1 or MG132 determined by flow cytometry. Statistical tests were carried out as in H. Data re shown as Mean±SD, n=3 independent experiments. **(J)** Volcano plot of the cell surface proteome of WT versus Rab7KO mouse fibroblasts identified by label-free MS. P-values are determined using two-sided permuted *t*-test with 250 randomizations. The black dotted line indicates the significance cutoff (FDR:0.05, S0:0.1) estimated by the Perseus software. n=3 biological replicates. Arbitrarily selected cell surface receptors are highlighted in red.

### Rab7KO destabilizes the cell surface proteome including Itgb1

To confirm that an inactivating mutation of Rab7 indeed decreases rather than increases Itgb1 surface levels, we disrupted the *Rab7a* gene (Rab7KO) in mouse kidney fibroblasts and the *RAB7A* gene in several human cell lines including HAP1 (myeloid), HEK (kidney), MCF7 (breast) and U2OS (bone) using the Crispr/Cas9 technology. Loss of Rab7 expression decreased total Itgb1 levels in lysates as well as on the cell surface (Figures 1D-E, S1A-C) of these cells, suggesting that stabilization of Itgb1 by Rab7 is a general principle of mammalian cells. The internalization kinetics of Itgb1 was unaffected by the loss of Rab7 expression in mouse fibroblasts (Figure S1D), whereas the degradation kinetics of the surface Itgb1 (Figure S1E) and the total Itgb1 (Figure S1F) was enhanced in Rab7KO cells. In line with the reduced Itgb1 surface levels, adhesion, spreading and proliferation were impaired in Rab7KO cells (Figure S1G-I).

The decreased Itgb1 levels in Rab7KO cells were efficiently rescued upon retrovirus-mediated re-expression of the EGFP-tagged Rab7 (Figure 1D-F). Furthermore, cells treated with Bafilomycin A1 (BafA1), which inhibits lysosomal protein degradation stabilized Itgb1, whereas MG132, which blocks proteasomal protein degradation was without effect (Figure 1G-I), which suggests that Itgb1 is degraded by lysosomes or lysosome-like organelles upon loss of Rab7 expression.

To investigate whether cell surface proteins other than Itgb1 are also downregulated on Rab7KO cells, we biotinylated the cell surface proteome of parental wild-type (WT) and Rab7KO mouse fibroblasts, precipitated the biotinylated proteins and compared their abundance by quantitative mass spectrometry (MS). The experiments revealed that Rab7KO cells displayed diminished surface levels of integrin family members (Itgb3, Itgb5, Itga2, Itga3, Itga5, Itga6 and Itgav) and numerous, integrin-unrelated proteins with different transmembrane topologies such as: amyloid-beta precursor protein (App), epidermal growth factor receptor (Egfr), low density lipoprotein receptor-related protein 1 (Lrp1), which are type I single-pass transmembrane proteins; transferrin receptor (Tfrc), a type II transmembrane protein and a classical marker of recycling endosomes; Piezo-type mechanosensitive ion channel component 1 (Piezo1), a multipass ion channel receptor; and Glypican-6 (Gpc6), a glycosylphosphatidylinositol(GPI)-anchored protein (Figure 1J). These results indicate that Rab7 loss impairs a general and not integrin-specific trafficking route.

### Rab7KO cells generate late endosomes and lysosomes

Our findings so far indicate that Itgb1 and numerous additional transmembrane proteins are decreased in the absence of Rab7 expression. This decrease requires functional lysosomes or lysosome-like organelles that express the integral proteins lysosomal-associated membrane protein 1 and 2 (Lamp1 and Lamp2), contain luminal lysosomal proteases such as the cathepsins that degrade substrates, establish an acidic luminal pH required for the function of lysosomal proteases, deliver substrates into the lumen for degradation and adopt a characteristic fingerprint-shaped ultrastructural morphology observed in the electron microscope (EM) as electron-dense membrane whorls (Huotari & Helenius, 2011; Saftig & Klumperman, 2009).

First, we immuno-stained Itgb1 and different lysosomal markers to reveal the subcellular localization in WT and Rab7KO fibroblasts (Figure 2A, S2A-C). In WT cells, Itgb1 was mainly observed in focal adhesions (FAs), ER and rarely in lysosomes. In Rab7KO cells, however, Itgb1 massively accumulated in lysosome-like structures that colocalized with Lamp1, Lamp2 and cathepsins such as cathepsin D (Ctsd), cathepsin B (Ctsb) and cathepsin L (Ctsl). An increased Pearson’s correlation coefficient (PCC) of Itgb1 with Ctsd (Figure 2B) and Itgb1 with Lamp1 (Figure S2D) confirmed their significant colocalization in Rab7KO fibroblasts.

**Figure 2.**
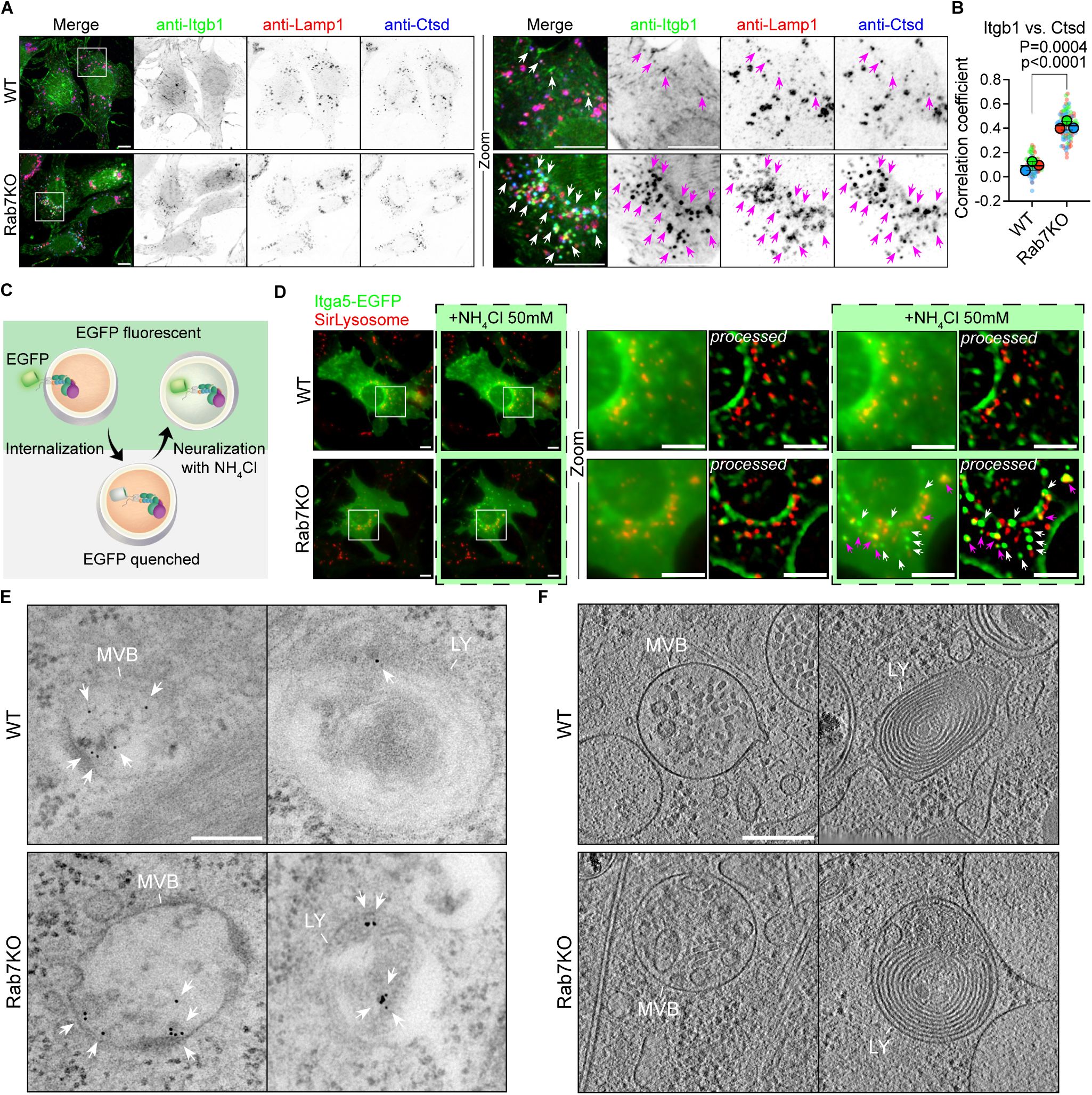
Itgb1 is delivered to *bona fide* lysosomes in Rab7KO cells. **(A)** Representative IF images of Itgb1, Lamp1 and Cathepsin D (Ctsd) in mouse fibroblasts. Arrowheads indicate Itgb1 accumulation in Lamp1- and Ctsd-positive lysosomes. Boxes indicate cell areas shown magnified in the Zoom panel. Sum intensity projections of confocal stacks are shown. Scale bar, 10µm. **(B)** Superplots showing Pearson correlation coefficient (PCC) between Itgb1 Ctsd in WT and Rab7KO mouse fibroblast. “P” indicates the p-value obtained by two-sided Welch’s *t*-test from the mean values of each independent experiment, N=3. “p” indicated the p-value obtained by two-sided Welch’s *t*-test from all individual values collected, WT n=82, KO n=103 cells. Bars represent Mean±SD of the mean values. **(C)** Schematic overview of EGFP-tagged Itga5 on the limiting membrane and in the lumen of lysosomes. The fluorophore signal is quenched by the intraluminal acidic pH and recovered upon neutralization with NH_4_Cl. **(D)** Representative widefield live-cell images of EGFP-tagged Itga5 and Sir^+^ lysosomes in mouse fibroblast before and after neutralization by NH4Cl. Boxes indicate cell areas shown in Zoom. Denoised images without and with (labeled as “processed”) background subtraction are shown. Arrowheads indicate Itga5-Itgb1 heterodimers in the acidic lumen of late endosomes (white arrowheads) and lysosomes (magenta arrowheads). Scale bar, 10µm. n≥3 independent experiments for each cell line. **(E)** Representative TEM images showing immunogold-labelled Itgb1 (arrowheads) in MVBs/late endosomes and lysosomes (LY). Scale bar, 0.2 µm. n=2 independent experiments with at least 2 EM grids imaged for each experiment. **(F)** Representative cryo-ET images showing morphology of MVBs/late endosomes and lysosomes (LY) in vitrified mouse fibroblast. Images show one slice of an electron tomography stack. Cells were stained with LysoTracker before vitrification to localize acidic organelles in cryo-fluorescence microscopy. Lamellae with a thickness around 100 nm were milled in the LysoTracker signal-rich area using focused ion beam (FIB). Scale bar, 0.2 µm. n=2 independent experiments with at least 2 different EM grids imaged for each experiment.

To determine whether the Itgb1 remains at the limiting membrane or is delivered into the lumen of Ctsd^+^ lysosomes, we tagged the cytoplasmic domain of the Itga5 subunit with the EGFP, whose fluorescent signal is pH-sensitive and quenched by ∼50% when the α5β1 integrin heterodimer is present in an acidified environment that is below pKa ∼5.5-6 (Figure 2C). We chose to EGFP-tag Itga5 to avoid modification of the cytosolic tail of Itgb1, which is important for integrin activation and trafficking (Bottcher et al., 2012; Rognoni et al., 2016; Steinberg et al., 2012). Itga5-EGFP signals were found on the cell surface and in a few endosomes of both, WT and Rab7KO cells, but were almost absent from lysosomes labeled with the Ctsd sensor Sir-Lysosome (Figure 2D). Since quenched EGFP can be recovered by neutralizing the pH of cells and lysosomes with NH_4_Cl, we treated WT and Rab7KO cells with NH_4_Cl. Whereas only marginal changes in Itga5-EGFP fluorescence intensity were observed in NH_4_Cl-treated WT cells, numerous bright Itga5-EGFP puncta were recovered in Rab7KO cells, with some being labelled with and some without Sir-Lysosome (Figure 2D, S2E-G). This finding indicates that acidified Sir-Lysosome^-^ late endosomes and acidified Sir-Lysosome^+^ lysosomes are present in Rab7KO cells and contain α5β1 integrin in their acidic lumen. Consistent with our immunofluorescence (IF) data, the dramatically increased amounts of luminal Itga5-contaning late endosomes and lysosomes suggest an increased lysosome targeting of integrins in Rab7KO cells, in comparison to WT cells (Figure S2F-G).

Finally, we used transmission EM to delineate the morphology of the Itgb1-containing endosomes in Rab7KO cells. To this end, we labeled cell surface Itgb1 with immunogold-coupled anti-Itgb1 antibodies and allowed the integrin-antibody complex to internalize for 2 hours. In both, WT as well as Rab7KO cells, the integrin-antibody complex localized to MVBs/late endosomes structures containing multiple small single membrane vesicles in their lumen, mature lysosomes filled with membrane whorls, and hybrid organelles formed upon fusion of lysosomes with MVBs/late endosomes (resulting in endolysosomes) and autophagosomes (resulting in autolysosomes), respectively (Figure 2E and S3A). Correlative light microscopy and cryo-electron tomography, which allow visualizing the morphology of endosomes and lysosomes in their native state, also revealed the presence of comparable endosomal and lysosomal morphologies in WT and Rab7KO cells (Figure 2F, S3B, Video S1). These findings indicate that the biogenesis of lysosomes and the intraluminal delivery of substrates takes place in the absence of Rab7.

### The Rab7KO lysosomes exhibit normal functions

Next, we tested whether the Rab7KO lysosomes exhibit genuine functions such as the degradation of autophagic materials and the exocytosis of the lysosomal content upon fusion with the plasma membrane. Functional lysosomes play a crucial role in macroautophagy (hereafter autophagy), a process that collects cytoplasmic materials and then delivers them to lysosomes for degradation (Mizushima, 2018; Yim & Mizushima, 2020). Autophagosomes, which are unable to mature into lysosomes on their own, were shown to fuse with pre-existing late-endosomes or lysosomes in a Rab7-dependent manner (Yim & Mizushima, 2020). Interestingly, recent studies challenged these findings and suggested that Rab7 is dispensable for autophagosome-lysosome fusion but required for the degradation of the autophagosome contents under fed, however, not under starved conditions (Kuchitsu et al., 2018). To examine whether autophagosomes fuse with Rab7KO lysosomes in our cell model, we performed the LC3-II flux assay (Klionsky et al., 2021). The cytosolic LC3-I (abbreviated for microtubule-associated protein 1A/1B-light chain 3; MAP1LC3), which serves as specific marker of autophagosome formation, is lipidated to LC3-II, incorporated into the inner and outer membrane of autophagosomes and eventually degraded upon fusion with lysosomes. Hence, the LC3-II levels report the LC3-I to LC3-II transition and the LC3-II degradation, of which the latter can be blocked by BafA1. Using our cell model, we found that in serum-cultured WT as well as Rab7KO fibroblasts, BafA1 treatment increased the LC3-II levels (Figure 3A-B), indicating that Rab7 is dispensable for basal autophagy, autophagosome-lysosome fusion, and degradation of autophagosomal proteins. Nutrient starvation further increased LC3-II levels in BafA1-treated WT and Rab7KO cells, which suggests that the response to an autophagic signal proceeds normally in the absence of Rab7, irrespective whether cells are fed or starved (Figure 3A-B).

**Figure 3.**
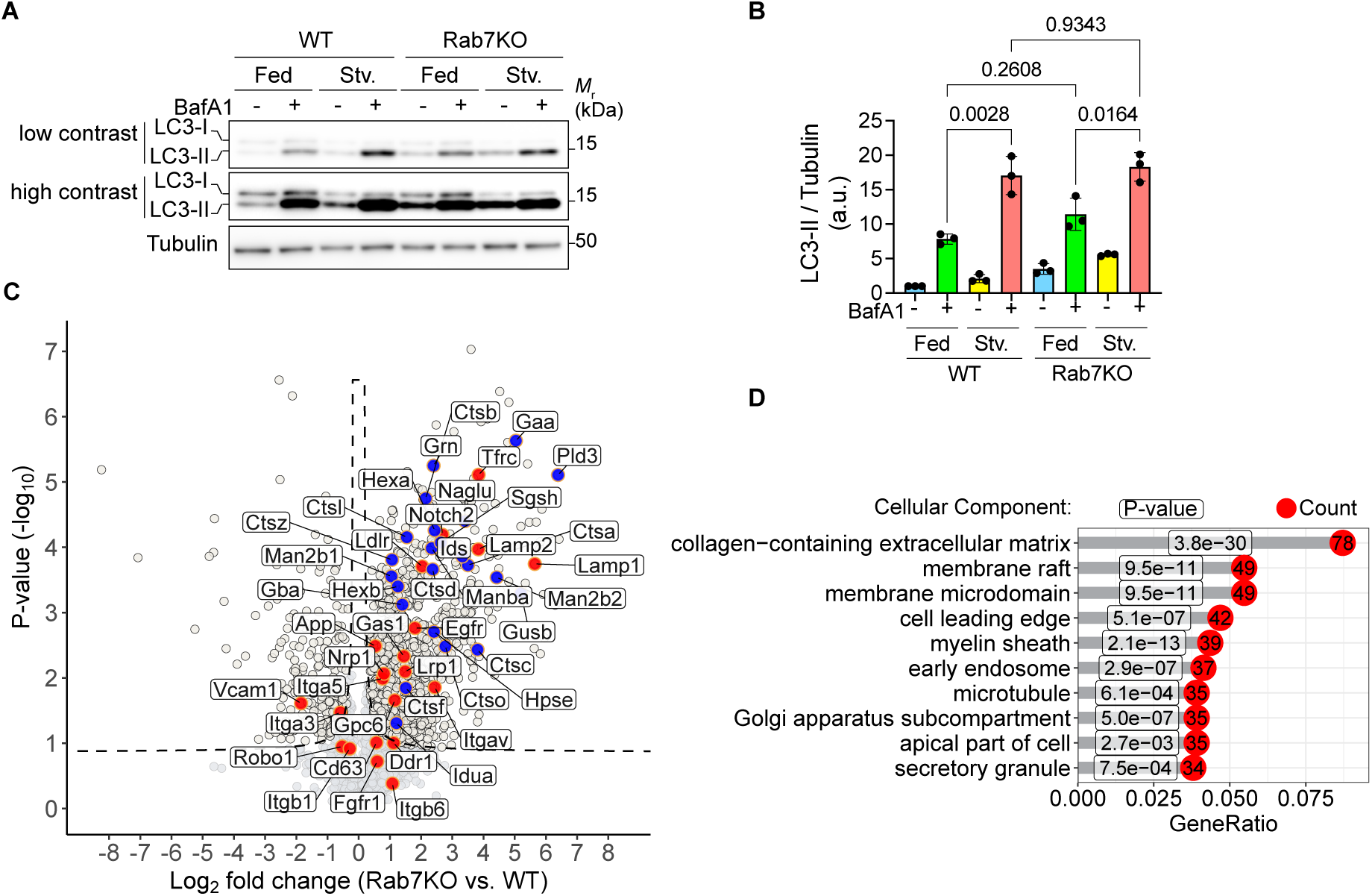
Rab7KO lysosomes function normally. **(A, B)** WB (A) and quantification (B) of LC3 in WT and Rab7KO mouse fibroblasts treated with and without BafA1. Tubulin served as loading control. Statistics was carried out by one-way ANOVA with Šidák’s post hoc tests. Data are shown as Mean±SD, n=3 independent experiments. **(C)** Volcano plot of the secretome of WT versus Rab7KO mouse fibroblast. The black dotted line indicates the significance cutoff (FDR:0.05, S0:0.1) estimated by the Perseus software. n=3 biological replicates. The arbitrarily highlighted cell surface receptors are indicated in red and representative lysosomal proteins in blue. **(D)** Gene ontology (GO) enrichment analysis of proteins showing a significant increase in the Rab7KO secretome. The top 10 GO terms regarding cellular components are displayed. P-values are show for each GO term and adjusted by the Benjamini-Hochberg (BH) method for controlling the FDR. Counts represent the number of genes found in the GO term. GeneRatio represents the ratio between the number of genes found in the GO term over total number of genes subjected to analysis.

In addition to degrading transmembrane proteins intracellularly, lysosomes and late endosomes also fuse with the plasma membrane and exocytose their luminal contents (Blott & Griffiths, 2002; Machado et al., 2021; Samie & Xu, 2014). To investigate whether Rab7KO lysosomes secrete their elevated cell surface proteome content, we measured the total secretome of serum-starved WT and Rab7KO mouse fibroblasts by MS. The experiment revealed that peptides from numerous cell surface proteins including integrins (Figure 3C; redish dots) and lysosomal enzymes such as Ctsb, Ctsd, Ctsl, Gaa, Grn, Pld3 were elevated in the Rab7KO secretome (Figure 3C; blue dots). To obtain a general overview of molecular functions affected by the activities of the secretome components, we performed a gene ontology (GO) enrichment analysis which revealed a significant over-representation of plasma membrane proteins with GO terms “membrane raft”, “membrane microdomain”, “cell leading edge”, “myelin sheath” and “apical part of cell” (Figure 3D). These data confirm an increased targeting of transmembrane proteins to lysosomes, and moreover, indicate that Rab7KO cells generate mature, functional lysosomes that secure protein degradation, autophagy progression and secretion of lysosomal contents.

### Rab7KO lysosomes are generated from recycling endosomes

Although our findings demonstrate that Rab7KO cells generate lysosomes with genuine lysosomal properties, our experiments so far do not answer as to why cell surface proteins are increasingly targeted to lysosomes in Rab7KO cells. The identification of the membrane source for Rab7KO lysosomes and the mechanism of Itgb1 delivery to Rab7KO lysosomes is key to understand the difference between the canonical, Rab7-dependent and the Rab7-independent lysosome biogenesis pathway. A first clue to these questions came from the cell surface proteome analysis of Rab7KO cells, which revealed in addition to the decreased Itgb1 levels, a dramatic reduction of Tfrc (Figure 1J), which is generally used as marker for recycling endosomes in WT cells. The decreased Tfrc surface levels were confirmed by flow cytometry and restored upon Rab7 expression (Figure S4A). Since in WT cells, Tfrc is barely sorted into late endosomes and lysosomes (Maxfield & McGraw, 2004), the reduced Tfrc surface levels in Rab7KO cells pointed to a malfunction of the recycling endosome pathway and increased degradation. This hypothesis was supported by immunostaining, which demonstrated co-staining of Tfrc and Itgb1 in Ctsd^+^ lysosomes and an increased PCC of Tfrc with Ctsd in Rab7KO compared to WT cells (Figure 4A-B).

**Figure 4.**
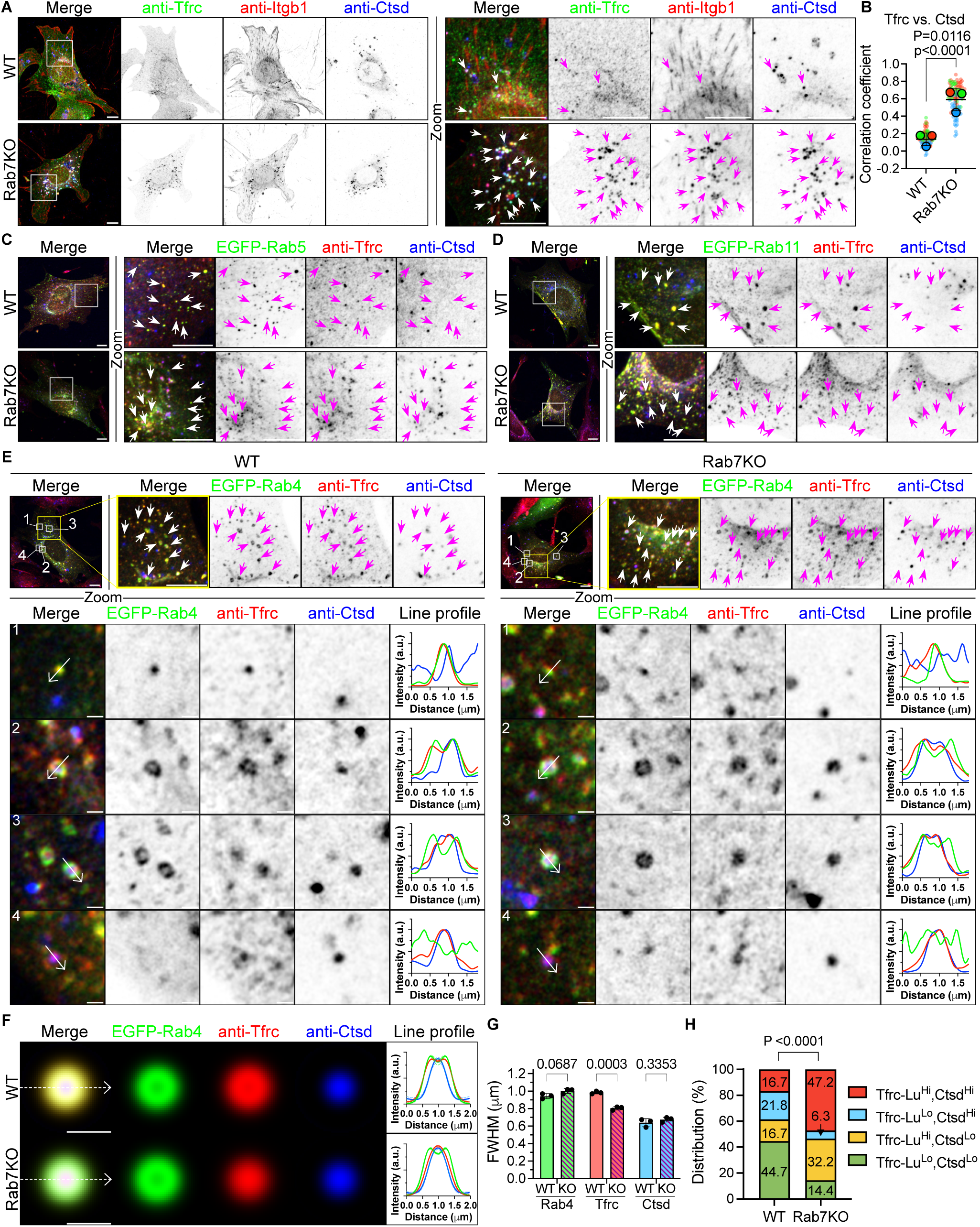
Rab7KO cells generate lysosomes from Rab4^+^ recycling endosomes. **(A)** Representative IF images of Transferrin receptor (Tfrc), Itgb1 and Ctsd in WT and Rab7KO mouse fibroblasts. Arrowheads indicate intracellular accumulation of Tfrc and Itgb1 in Ctsd-positive lysosomes. Boxes indicate cytoplasmic areas shown in Zoom. Sum intensity projections of confocal stacks are shown. Scale bar, 10µm. **(B)** Superplots showing PCC between Tfrc and Ctsd in WT and Rab7KO mouse fibroblast. “P” indicated the p-value obtained by two-sided Welch’s *t*-test from the mean values of each independent experiment, N=3. “p” indicated the p-value obtained by two-sided Welch’s *t*-test from all individual values collected, WT n=100, KO n=120 cells. Bars represent Mean±SD of the mean values. **(C, D)** Representative IF images of Tfrc and Ctsd in WT and Rab7KO mouse fibroblasts expressing EGFP-Rab5 (C) and EGFP-Rab11 (D), respectively. Arrowheads indicate colocalization of Rabs and Tfrc. Boxes indicate cytoplasmic areas shown at in Zoom. Sum intensity projections of confocal stacks are shown. Scale bar, 10µm. **(E)** Representative IF images of Tfrc and Ctsd in WT and Rab7KO mouse fibroblasts expressing EGFP-Rab4. Arrowheads indicate colocalization of Rab4 and Tfrc. Yellow boxes indicate cytoplasmic areas shown in Zoom. Numbered white boxes indicate endosomes of different classes. Arrows indicate the direction of line profiles of EGFP-Rab4 (green), Tfrc (red) and Ctsd (blue). The upper panels show sum intensity projections of confocal stacks. The lower panels show a single confocal slice. Each line profile was produced from a single slice. Scale bar: upper, 10µm; lower, 1µm **(F)** Image of modelled Rab4^+^ endosomes in WT and Rab7KO cells. Rab4^+^ endosomes with a donut-like shape (n=516 from WT and n=505 from Rab7KO cells) were collected and normalized in intensity and isotropicity from three independent experiments and in total of 62 WT and 58 Rab7KO cells expressing EGFP-Rab4. Arrows indicate the direction of the line profiles. Dashed lines show the line profiles of each independent experiment. The line profile generated from the input of all cells is shown as solid lines in the right panel. **(G)** Quantification of the full width at half maximum (FWHM) of Rab4, Tfrc and Ctsd line profiles generated in the independent experiments. Statistics was calculated by the Two-sided Welch’s *t*-test. Data are shown as Mean±SD, n=3 independent experiments. **(H)** Classification of Rab4^+^ endosomes based on the degree of luminal levels of Tfrc (Tfrc-Lu) and Ctsd (Ctsd), categorized into high (Hi) or low (Lo) levels. P-value for the contingency test is determined using Chi-squared test. See also Figure S4E-F.

The increased lysosomal targeting of Tfrc suggests that recycling endosomes play a role in the formation of Rab7KO lysosomes. To test this hypothesis, we expressed EGFP-tagged forms of the three major Rab proteins orchestrating Tfrc recycling (Sonnichsen et al., 2000): Rab4, which is associated with (classical) recycling endosomes; Rab5, which is associated with endocytic vesicles and early endosomes that can, to a low extent, directly recycle surface proteins back to the plasma membrane; and Rab11, which is associated with recycling endosomes and the PNRC.

Our immunostaining revealed that Tfrc colocalized with Rab4, Rab5 and Rab11 on endosomal vesicles in WT as well as Rab7KO cells (Figure 4C-E), indicating that Tfrc is present in all three trafficking routes irrespective whether Rab7 is expressed or not. Since the overexpressed Rab5 and Rab11 resulted in the colocalization of Tfrc with Rab5 or Rab11, however, not with Ctsd in WT as well as Rab7KO cells (Figure 4C-D), we conclude that Tfrc^+^/Ctsd^+^ lysosomes are neither directly generated from Rab5^+^ nor from Rab11^+^ endosomes.

Overexpression of Rab4 resulted in colocalization of Tfrc and Rab4 on Ctsd^-^ and Ctsd^+^ structures of different sizes, ranging from puncta at the limit of optical resolution to large vesicular structures with a diameter up to 1µm in WT and Rab7KO cells (Figure 4E). A thorough examination of the vesicular structures revealed that four different classes of organelles could be distinguished both, in WT and Rab7KO cells, which differed in size, in Rab4 and Ctsd signals and in the localization of Tfrc on the limiting membrane or in the lumen (Figure 4E, box 1-4). One class of small structures showed colocalization of Tfrc and Rab4 and absence of Ctsd, which points to classical recycling endosomes (see box 1). A second class of large vesicles showed Rab4 and Tfrc colocalizing at the limiting membrane and Ctsd in the lumen both, in WT and Rab7KO cells, indicating that these structures originate upon fusion of Rab4^+^ recycling endosomes with Ctsd^+^ carriers (see box 2). A third class also of large vesicles showed Rab4 at the limiting membrane and Tfrc as well as Ctsd in the lumen indicating endosomal maturation with internalized, luminal Tfrc (see box 3). Finally, a fourth of small structures lacked Rab4 but was positive for Tfrc and Ctsd signals, indicating that Rab4 dissociated from the endosomal membrane during lysosome conversion (see box 4). The unconventional Tfrc^+^/Rab4^+^/Ctsd^+^ endo/lysosomal organelles (class 2 and 3) resemble late endosomes/endolysosomes that are likely formed by the fusion of recycling endosomes with carriers containing lysosomal enzymes.

### Rab4^+^ late endosomes and Rab4^+^ endolysosomes generate lysosomes

Although Rab4^+^ late endosomes and Rab4^+^ endolysosomes were apparent to a much lesser extent in WT compared to Rab7KO cells (Figure 4E), we asked how we can most accurately determine their numbers in WT and Rab7 cells. The question could be addressed by determining the census of the four vesicular structures in WT and Rab7KO cells. The census determined by immunostaining, however, would produce inaccurate numbers as the transition of recycling endosomes to late endosomes and finally lysosomes is a continuous, highly dynamic process with intermediate structures in which immunosignals will be low and therefore difficult to flawlessly detect and assign to a specific class of vesicular structures. For example, Tfrc can be present at the limiting membrane as well as in the lumen at the same time, or Ctsd may initially be present at a low level that is difficult to distinguish from background noise and gradually increase during the transition to late endosomes and lysosomes.

To overcome this hurdle, we developed an image-based flux-like assay focusing on the large class 2 and class 3 donut-like structures whose limiting membrane and lumen can be optically resolved (Figure S4B). The measurement of Rab4, Tfrc and Ctsd immunosignals in these structures was used to deduce a quantitative transition from Tfrc^+^/Rab4^+^/Ctsd^-^ recycling endosomes to Tfrc^+^/Rab4^+^/Ctsd^+^ endo/lysosomal organelles. In our assay we compared the full <u>w</u>idth of a fluorescence line profile at <u>h</u>alf <u>m</u>aximum (FWHM), which is a statistical parameter used to describe the width of a function. In our case, the function described by FWHM is a Gaussian-like distribution of fluorescence signals and represents the distance between points on the Gaussian curve (width) at half of the maximum value. Fluorescence signals emitted from the endosome lumen generate a function of a single Gaussian-like distribution (SGD), whereas fluorescence emitted at the limiting membrane generate a function of a double Gaussian-like distribution (DGD). The FWHM measurements of Tfrc and Ctsd on hundreds of class 2 and class 3 structures allows to accurately measure their size (which differs if the signal is emitted at the limiting membrane or in the lumen), the flux of membrane proteins such as Tfrc from the limiting membrane into the lumen, and the gradual accumulation of lysosomal proteases occurring in the lumen during lysosome maturation.

To obtain the quantitative FWHM assessment of Tfrc, we normalized the intensity and isotropicity of the Tfrc signal of donut-like endosomes in WT and Rab7KO cells, respectively, and then made an average projection to create a model endosome (Figure S4B). Since Rab4 should localize to the outer leaflet of the limiting membrane, the Rab4 signal should produce DGD profiles and the dimension of the FWHM should reflect the size of the endosome. The Ctsd signal, on the other hand, should produce SGD profiles and only be emitted from inside the lumen. Accordingly, Rab4 produced a DGD profile and an average FWHM of about 1μm and Ctsd SGD profiles and a FWHM of 0.6μm in WT as well as Rab7KO cells (Figure 4F-G), indicating that the size of the large, donut-like endosomes is similar in both cell lines. The Tfrc also produced DGD profiles in WT cells with a FWHM, similarly like for Rab4, of around 1μm, indicating that in WT cells the majority of Tfrc colocalizes with Rab4 at the limiting membrane. In sharp contrast, in Rab7KO cells Tfrc produced SGD profiles with a dramatically decreased FWHM, indicating that the majority of Tfrc was internalized from the limiting membrane into the lumen. We also observed that the accumulation of Ctsd in donut-like endosomes, calculated as ratio of Ctsd intensity in endosomes versus Ctsd intensity in whole cells, was increased in Rab7KO compared to WT cells (Figure S4C). In line with this finding, also the PCC revealed that the correlation of Tfrc and Ctsd immunosignal in co-staining experiments was significantly higher in Rab7KO compared to WT cells (Figure S4D). The underlying reason for the very low Tfrc and Ctsd flux in WT compared to Rab7KO cells is very likely due to a diminished ability of recycling endosomes to mature into late endosomes and lysosomes when the Rab7-mediated conventional lysosomal maturation pathway prevails.

Finally, we measured the ratio of the FWHM of Tfrc versus Rab4 in individual endosomes with either high or low Ctsd levels to obtain a semiquantitative measure of the endo/lysosome maturation. A high Tfrc-over-Rab4 ratio indicates less internalization and thus low luminal Tfrc, whereas a low Tfrc-over-Rab4 ratio indicates more internalization and thus high luminal Tfrc in Ctsd^high^ and Ctsd^low^ endosomes. Hence, (1) low luminal Tfrc and Ctsd indicate classical recycling endosomes, (2) either high luminal Tfrc or high Ctsd indicates maturing intermediates between recycling endosomes and endolysosomes, and (3) high luminal Tfrc as well as high luminal Ctsd indicate endolysosomes. The experiment revealed that in WT cells, 44.7% of Rab4^+^ endosomes showed low luminal levels of Tfrc and Ctsd and were classified as classical recycling endosomes, 38.6% displayed high levels of either Tfrc or Ctsd and were classified as maturing endosomes and 16.7% showed high luminal levels of Tfrc as well as Ctsd and were classified as endolysosomes (Figure 4H, S4E-F). In Rab7KO cells, the distribution of the three classes of particles shifts from classical recycling endosomes to endolysosomes: 14.4% exhibited low luminal levels of Tfrc and Ctsd, 38.5% displayed high luminal signals of either Tfrc or Ctsd, and 47.2% showed high luminal levels of Tfrc as well as Ctsd. Altogether, these results imply that the generation and size control of Rab4^+^ enlarged endosomes are Rab7 independent and that loss of Rab7 promotes the maturation of Rab4^+^ recycling endosomes towards lysosomes.

### Biochemistry confirms systematic shift of lysosome biogenesis pathway

The cell imaging studies suggest that in Rab7KO cells the cell surface proteins are routed from recycling endosomes via Rab4^+^ late endosomes to lysosomes, which results in the massively decreased their surface levels. To confirm this finding biochemically, we expressed GFP-tagged Rab4 or Rab5 in WT and Rab7KO fibroblasts, immuno-isolated intact Rab4^+^ or Rab5^+^ endosomes using anti-GFP antibody-coupled beads and compared their protein contents using quantitative MS (Figure 5A). In WT cells, Rab5^+^/Rab7^+^ endosomes are the source of lysosome biogenesis and therefore lysosomal protein should be detected in Rab5^+^ endosomes but not in Rab4^+^ recycling endosomes. Indeed, we found that lysosomal proteins, including lysosomal membrane-associated proteins (e.g. the Lamtors/Regulator complex), transmembrane proteins (e.g. Lamp1 and Lamp2) and lysosomal luminal proteins (e.g. cathepsins) were enriched in Rab5^+^ but not Rab4^+^ endosomes of WT cells (Figure 5B). In sharp contrast, in Rab7KO cells these proteins were enriched in Rab4^+^ and not in Rab5^+^ endosomes (Figure 5C-D). This finding together with the GO ontology analysis showing specific enrichment of proteins from the classical late-endosome/lysosome pathway (GO term “late endosome”, “vacuolar membrane”, “lytic vacuole membrane” and “lysosomal membrane”; Figure 5E) for Rab4^+^ endosomes in Rab7KO cells independently confirms that lysosomes are primarily generated from Rab4^+^ rather than Rab5^+^/Rab7^-^ endosomes in Rab7KO cells (Figure 6).

**Figure 5.**
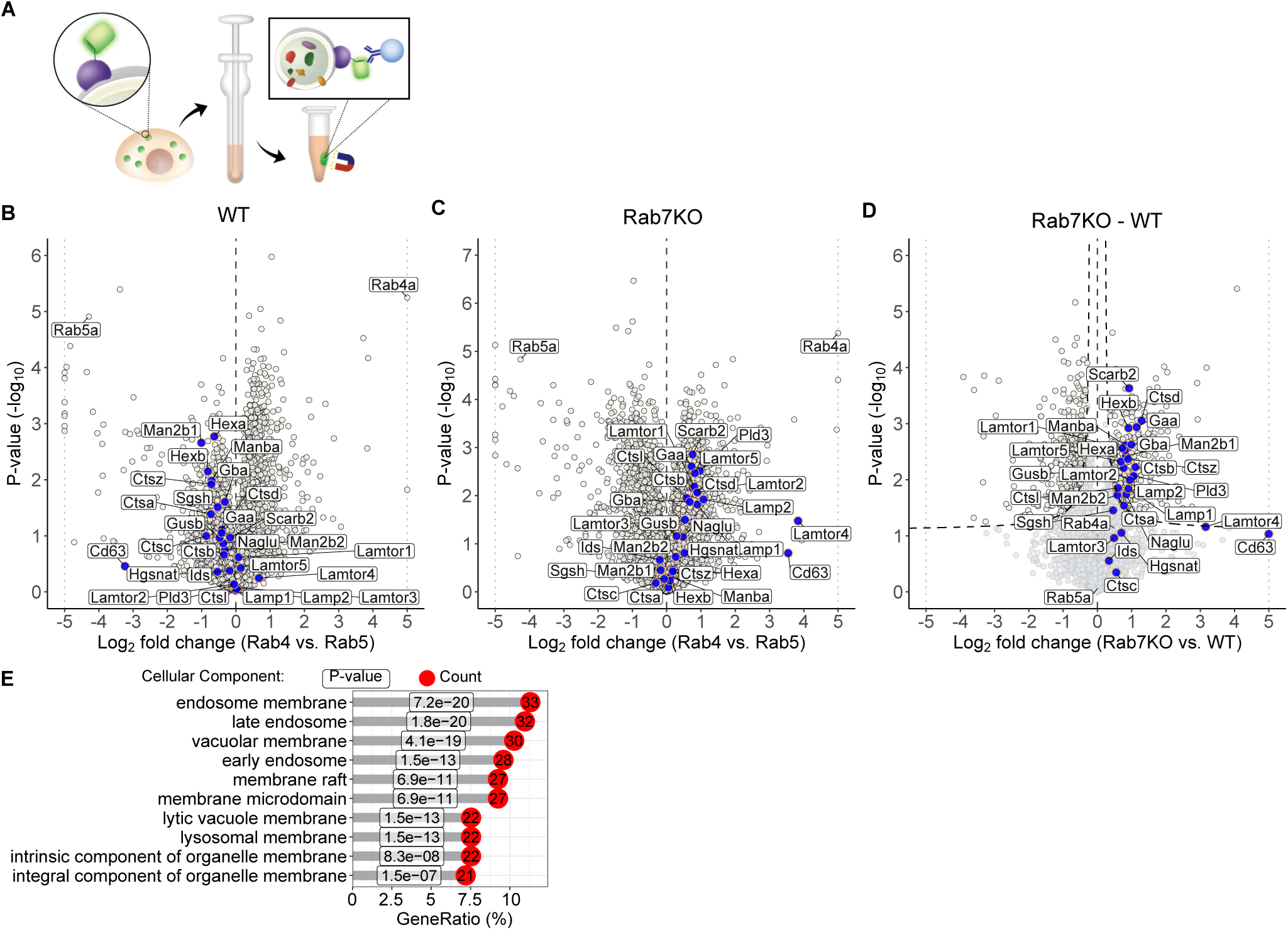
Rab4^+^ endosomes outcompete Rab5^+^ endosomes for lysosome biogenesis in Rab7KO cells. **(A)** Schematic representation showing the immunoisolation of intact EGFP-Rab4^+^ and EGFP-Rab5^+^ endosomes for proteomic profiling. **(B, C)** Volcano plot of proteins in Rab4^+^ versus Rab5^+^ endosomes of WT (B) and Rab7KO (C) cells. Representative lysosomal proteins are highlighted in blue. Out-of-range values (outside the x-axis range of −5 to 5) are plotted on the border. **(D)** Volcano plot showing the relative difference in proteins localization in Rab4^+^ endosomes in Rab7KO vs. WT cells. P-values are determined using two-sided permuted t-test with 250 randomizations. The black dotted line indicates the significance cutoff (FDR:0.05, S0:0.1) estimated by the Perseus software. n=3 biological replicates. **(E)** Gene ontology enrichment analysis of proteins with significant shifts from Rab5^+^ to Rab4^+^ endosomes. The top 10 GO terms are displayed. P-values are show for each GO term and adjusted by the Benjamini-Hochberg (BH) method for controlling the FDR. Count represents number of genes found in the GO term. GeneRatio represents ratio between number of genes found in the GO term over total number of genes subjected to analysis.

**Figure 6.**
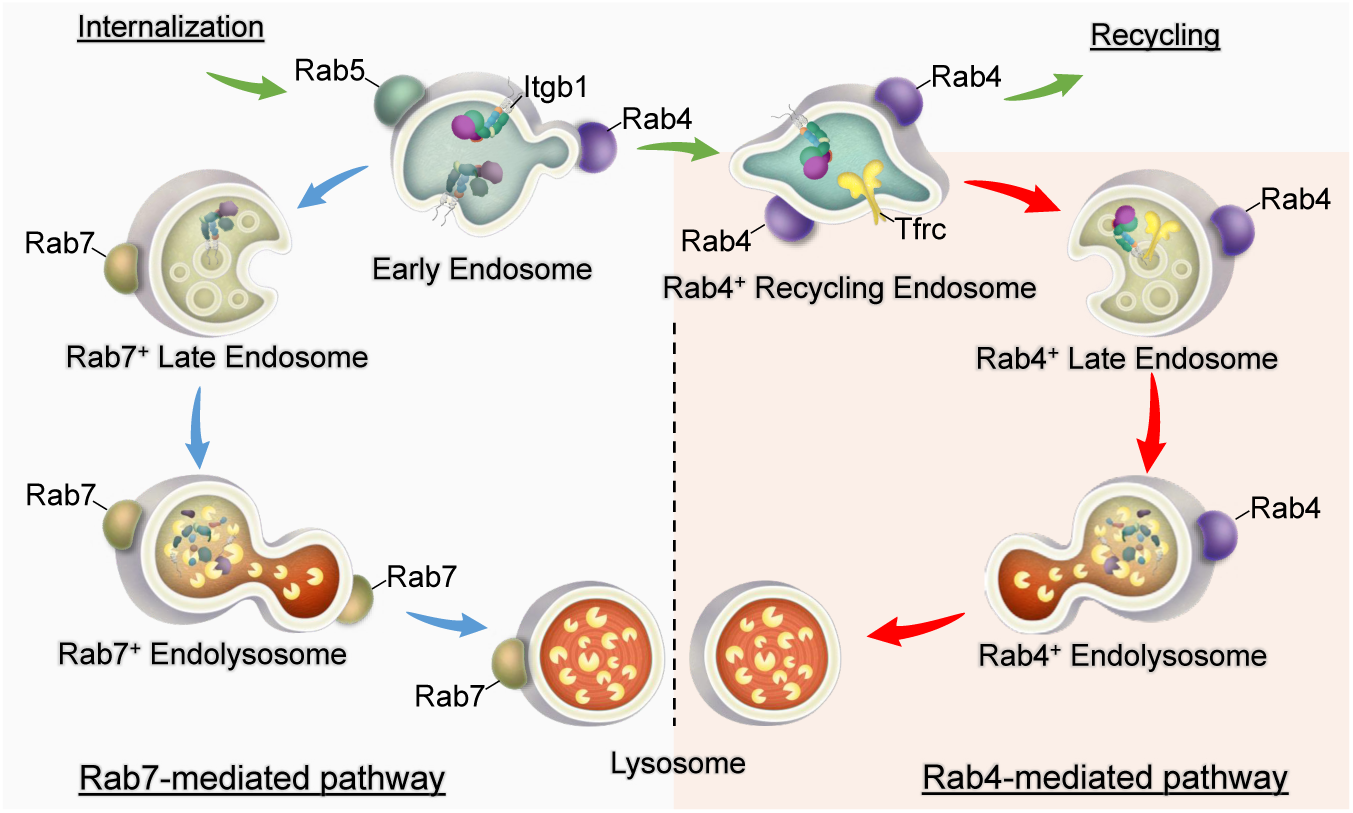
Schematic representation of integrin and Tfrc degradation via the Rab7- or Rab4-mediated lysosome biogenesis pathway. In the canonical Rab7-mediated lysosome biogenesis pathway, late endosomes mature into lysosomes by fusing with carriers containing lysosomal enzymes. In the Rab4-mediated non-canonical lysosome biogenesis pathway operating upon Rab7 loss, lysosomes are generated from Rab4^+^ recycling endosomes, resulting in the degradation of proteins originally routed to the recycling pathway.

### Rab4 overexpression generates endo/lysosomes and decreases protein surface levels in WT cells

Since expression of EGFP-Rab4 leads to the appearance of Rab4^+^ endo/lysosomal organelles in WT cells, although to a much lesser extent than in Rab7KO cells, we assessed Itgb1 and Tfrc surface levels in Rab4-overexpressed WT cells. The experiment revealed decreased Tfrc surface levels (that were similarly low like in EGFP-transfected Rab7KO cells) and only slightly decreased Itgb1 surface levels (Figure S4G-H). Interestingly, the Tfrc surface levels further decreased when Rab4 is overexpressed in Rab7KO cells, whereas Itgb1 levels did not significantly differ between Rab4- and EGFP-expressed Rab7KO cells (Figure S4G-H). These findings indicate that Rab4 overexpression generates late endosomes in WT cells, which receive large amounts of Tfrc and less Itgb1 from Rab4^+^ recycling endosomes.

Similarly like in mouse fibroblasts, the human MCF7 breast cancer cell line and the human U2OS sarcoma cell line also increased lysosomal targeting of Tfrc and Itgb1 upon deleting the *RAB7A* gene (Figure S5A-D and S6A-D), indicating the Rab4-mediated lysosomal pathway is also activated in these cells upon Rab7 loss. The WT and Rab7KO U2OS cells contained, in contrast to mouse fibroblasts and human MCF7 cells, particularly large Tfrc^+^ structures with the Tfrc signals enriched on the limiting membrane and Ctsd signals enriched in the lumen (Figure S6C), suggesting that Rab4^+^ recycling endosomes fused in WT and Rab7KO U2OS cells with Ctsd carriers without the need to manipulate the endogenous Rab4 levels. The overexpression of EGFP-Rab4 further increased the formation of Rab4^+^/Tfrc^+^/Ctsd^+^ structures in Rab7KO UO2S cells, induced these structures in Rab7KO MCF7 cells and to a lesser extent in WT MCF7 and U2OS cells (Figure S5E and S6E), which altogether demonstrates that the Rab4-mediated lysosome biogenesis pathway operates in mouse and human cells.

## Discussion

Our whole genome screen in HAP1 cells for regulators of Itgb1 surface stability identified Rab7 as unexpected candidate as stabilizer of Itgb1 and many additional cell surface proteins including Tfrc. Whereas genetic loss-of-function studies of Rab7 lead to embryonic lethality in all animal models tested (Cherry et al., 2013; Kawamura et al., 2012; Kreis et al., 2021; Roy et al., 2013; Skorobogata & Rocheleau, 2012), deficiency of Rab7 in mammalian cells (Kuchitsu et al., 2018; Roy et al., 2013; Schleinitz et al., 2023) produced viable and apparently normally appearing cells *ex vivo*, indicating that the developmental arrest *in vivo* must underlie severe defect(s) that are not obvious *ex vivo*. Since the massive degradation of surface proteins likely accounts for the embryonic lethality *in vivo*, we decided to investigate the mechanism that underlies this unexpected finding.

In search for a mechanistic explanation, we identified a novel lysosome maturation pathway, in which Rab4^+^/Tfrc^+^ recycling endosomes generate endo/lysosomal organelles (Figure 6). The Rab4^+^ late endosomes/endolysosomes, in the style of the canonical Rab7^+^ late endosomes/endolysosomes, acquire membrane proteins such as Lamp1 and Lamp2, contain hydrolytic enzymes such as the cathepsins, an acidic environment in their lumen, and further mature into lysosomes filled with characteristic membrane whorls (Huotari & Helenius, 2011; Saftig & Klumperman, 2009). In support of these findings, an orthogonal, biochemistry-based assay with Rab7KO cells also revealed that Rab4^+^ and not Rab5^+^ endosomes were enriched with late endosome- and lysosome-specific proteins including membrane associated proteins such as the LAMTOR/Ragulator complex (Laplante & Sabatini, 2009), transmembrane proteins such as Lamp1 and 2, and hydrolytic enzymes. Expectedly, in WT cells these proteins were enriched on Rab5^+^ and not Rab4^+^ endosomes. Furthermore, the Rab4-induced lysosomal pathway described and characterized here for Rab7-null mouse fibroblasts is also activated upon loss of Rab7 in all cell types that were analyzed in this study.

Our immunostainings showed that Tfrc^+^/Ctsd^+^ lysosomes are readily detected in several Rab7-deficient cell lines and only sporadically in WT cell lines. Upon expression of EGFP-Rab4, the abundance of Rab4^+^/Tfrc^+^/Ctsd^+^ endo/lysosomal organelles increased in Rab7KO and also became more obvious in the WT cell lines that we analyzed. Since the maturation of lysosomes from recycling endosomes is a fluent process with intermediate organelles that cannot be unequivocally assigned to either recycling endosomes, late endosomes or endolysosomes, it is difficult to accurately determine their census and compare the abundance of Rab4^+^ endo/lysosomal organelles between WT and Rab7KO cells. To overcome this obstacle, we determined size and sub-organelle localization of Rab4, Tfrc and Ctsd by measuring the full width at half maximum (FWHM) of their fluorescence signals. These measurements revealed a clear bias of endo/lysosome maturation characterized by an elevated accumulation of luminal Tfrc and Ctsd in Rab7KO compared to WT cells, while their size was similar between WT and Rab7KO cells. In light of the presence of Rab4^+^ endo/lysosomal organelles also in WT cells, it is conceivable that the Rab7- and the Rab4-induced protein degradation pathways can principally act in parallel, and that the latter may become relevant in cells which contain low Rab7 and/or high Rab4 levels. It will be important in future to experimentally define condition(s) in which the Rab4-induced protein degradation pathway is activated and outweighs the Rab7 pathway and degrades cell surface proteins that are actually designated to recycle back to the plasma membrane.

It is well known that Rab7 promotes the formation of late endosomes and their subsequent maturation into endolysosomes and further into lysosomes to ensure that proteins designated for degradation are degraded, and proteins designated for recycling to the plasma membrane are routed by the activity of Rab4 from early endosomes into recycling endosomes and are not degraded. The ability of Rab4^+^ recycling endosomes to principally route cargo to lysosomes for degradation, although to a very small scale, suggests that in WT cells Rab7 outcompetes Rab4 enabling lysosome fusion on the acceptor membrane. The competition between Rab7 and Rab4 might be based on the higher affinity of the Rab7 for recruiting proteins that regulate membrane fusion such as tethering complexes, SNAREs, SNARE regulators and additional small GTPases via Rab7 effectors or via the microenvironment of Rab7 microdomains (de Araujo et al., 2020; Guerra & Bucci, 2016; Langemeyer et al., 2018).

Our immunostaining also demonstrated that, in contrast to Rab4, neither overexpressed Rab5 nor Rab11 colocalized with lysosomal markers. Given that Rab4, Rab5 and Rab11 can be present on the same endosomes but in distinct microdomains (Sonnichsen et al., 2000), it remains to be shown whether Rab4^+^ endo/lysosomal organelles originate from endosomes solely decorated with Rab4 or from endosomes that also harbor Rab5 and Rab11 which become rapidly lost during maturation. The mechanism and timing of Rab4 dissociation from maturing lysosomes and the involvement of other small GTPases are important questions that need to be addressed in future studies.

We also found that autophagosome-lysosome fusion or plasma membrane-lysosome fusion are unaffected by the loss of Rab7 in our fibroblast model. The LC3-flux assay showed that LC3-II is delivered to both, WT and Rab7KO lysosomes for degradation irrespective whether cells are serum starved or serum treated, indicating that autophagosome-lysosome fusion does not require the activity of Rab7. These findings contradict the previous studies reporting that Rab7KO compromises autophagy (Kuchitsu et al., 2018; Roy et al., 2013). In both studies, however, serum starvation failed to increase LC3-II levels, *i.e.* induce autophagy which could be caused by the specific cell handling or the cell models used in these studies. Interestingly, the quantitative and qualitative measurements of the secretome also indicate that fusion of late endosomes and lysosomes with the plasma membrane proceeds in a Rab7-independent manner. Quantitatively, the Rab7KO cells released more cargo such as luminal enzymes and cell surface receptors than WT cells, which, however, is expected from the massive ‘misrouting’ of proteins normally destined to recycle into the lysosomal degradation pathway. Given the involvement of the lysosomal secretome in various physiological and pathological conditions (Blott & Griffiths, 2002; Buratta et al., 2020; Lee & Ye, 2018), switching the canonical, Rab7-directed or non-canonical, Rab4-directed protein degradation pathway, e.g. by decreasing Rab7 and/or increasing Rab4 levels, can have significant consequences not only due to the decrease of the surface proteome but also due to the abundant secretion of lysosomal contents.

## Acknowledgment

We thank the sequencing, mass spectrometry and imaging facilities of the Max Planck Institute of Biochemistry and the EM-Histo Lab of the Max Planck Institute for Biological Intelligence for the invaluable support. This work was supported by the European Research Council (ERC) under the European Union’s Horizon 2020 research and innovation program (grant agreement No. 810104 – Point) and the Max Planck Society.

## Author contribution

Conceptualization and writing: GW and RF; Investigation: GW, XP, KY and SG; Supervision and Funding Acquisition: RF.

---

**Figure S1.**
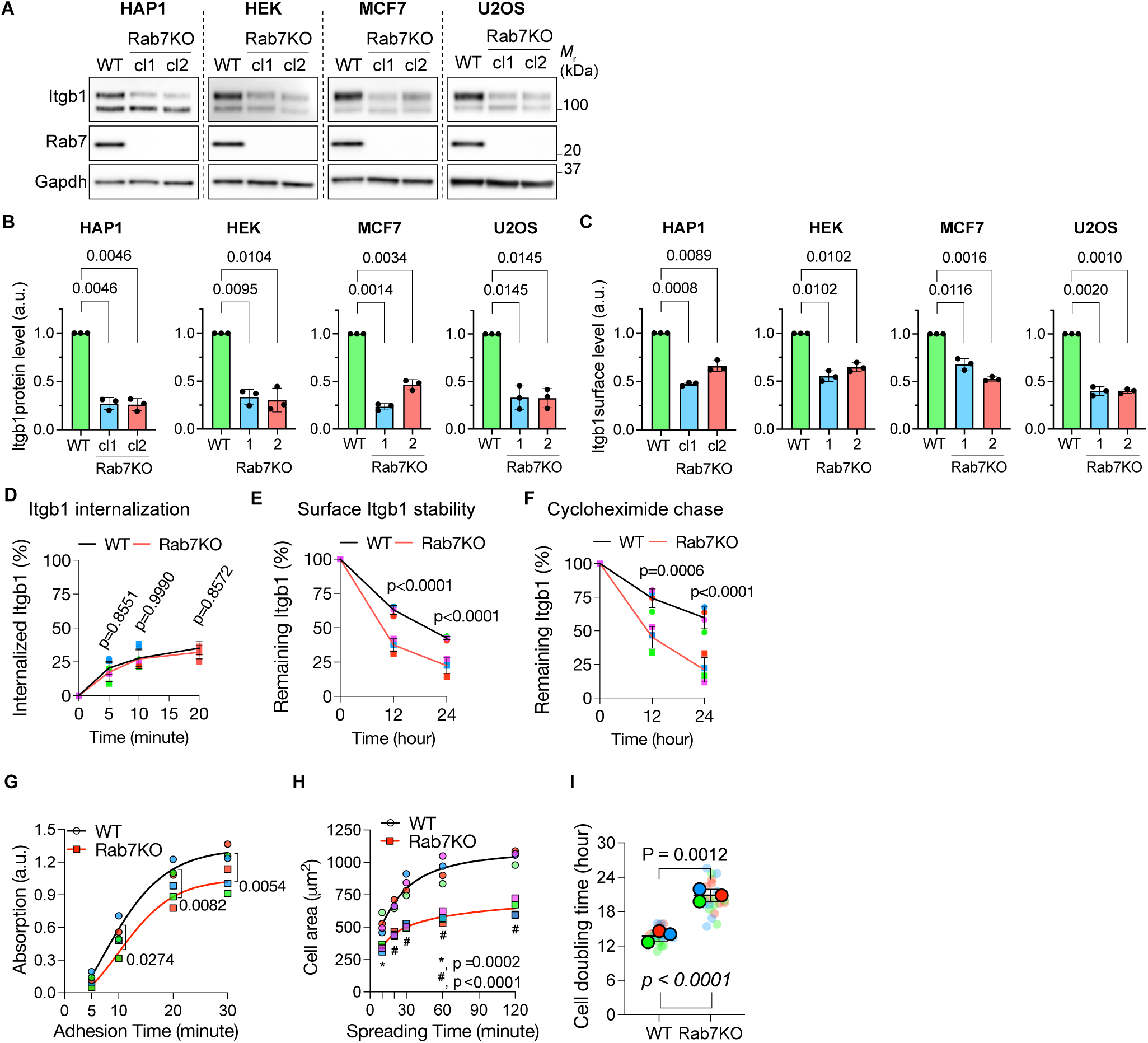
Related to Figure 1. **(A, B)** WB (A) and quantification (B) of Itgb1 in WT and Rab7-null clones derived from HAP1, HEK, MCF7 and U2OS cells. HAP1 cl.1 and cl.2 are independently generated Rab7KO clones. HEK, MCF7 and U2OS cell lines indicated as KO1 and KO2 are expanded pools derived from 100 flow cytometry sorted cells. The matured form of Itgb1 (upper band) was quantified in (B). Statistics was analyzed by two-sided multiple paired *t*-test with Holm-Šidák correction. Data are shown as Mean±SD, n=3. **(C)** Itgb1 surface levels on indicated cell lines determined by flow cytometry. Statistical tests were carried out as in B. Data are shown as Mean±SD, n=3 independent experiments. **(D)** Quantification of Itgb1 internalization kinetics in WT and Rab7KO mouse fibroblasts. Biotinylated proteins were pulled down by streptavidin beads and the amount of Itgb1 were measured by capture-ELISA. Statistics was analyzed by two-way ANOVA with Šidák’s post hoc tests. Mean±SD, n=4 independent experiments. **(E)** Quantification of surface Itgb1 degradation kinetics in WT and Rab7KO mouse fibroblasts. The amount of Itgb1 remaining over indicated times were measured by capture-ELISA. Statistics was analyzed by two-way ANOVA with Šidák’s post hoc tests. Data are shown as Mean±SD, n=4 independent experiments. **(F)** Quantification of total Itgb1 degradation kinetics in WT and Rab7KO mouse fibroblasts using the cycloheximide chase assay. Cells were lysed at indicated times and Itgb1 levels were measured by WB. Statistics was analyzed by two-way ANOVA with Šidák’s post hoc tests. Data is shown as Mean±SD, n=4 independent experiments. **(G)** Numbers of adherent WT and Rab7KO mouse fibroblasts at indicated time points after seeding on FN-coated glass. Symbols represent mean values of independent experiments; lines sigmoidal curve fit and numbers indicate p-values determined by two-way ANOVA with Šidák’s post hoc tests, n=4 independent experiments. **(H)** Cell spreading area of WT and Rab7KO mouse fibroblasts on FN-coated glass surface at indicated time points after seeding. Colored symbols represent mean values of independent experiment; lines represent sigmoidal curve fit; numbers p-values determined by two-way ANOVA with Šidák’s post hoc tests, n=4 independent experiments. **(I)** Superplots showing cell doubling time of WT and Rab7KO mouse fibroblasts. “P” indicated the p-value obtained by two-sided Welch’s *t*-test from the mean values of each independent experiment, N=3. “p” indicated the p-value obtained by two-sided Welch’s *t*-test from all individual data points collected, n=24 imaged areas. Bars represent the Mean±SD of the mean values.

**Figure S2.**
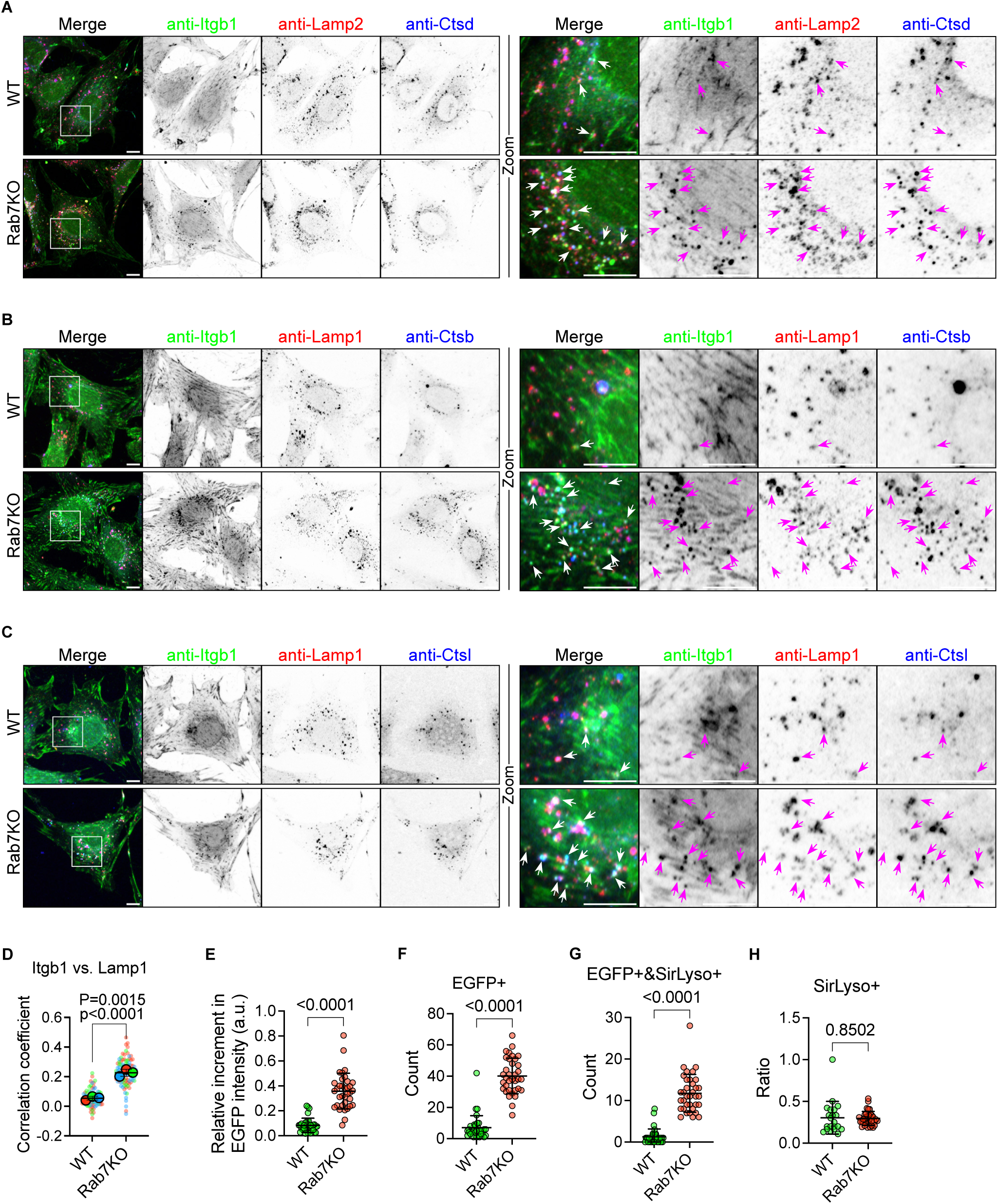
Related to Figure 2. **(A-C)** Representative confocal image sections of WT and Rab7KO mouse fibroblasts. Cells were immunostained for Itgb1, Lamp2 and Ctsd in (A); Itgb1, Lamp1 and Cathepsin B (Ctsb) in (B); and Itgb1, Lamp1 and Cathepsin L (Ctsl) in (C). Arrowheads indicate triple colocalization of Itgb1, Lamp and Cathepsin. Boxes indicate cytoplasmic areas shown in Zoom. Sum intensity projections of confocal stacks are shown. Scale bar, 10µm. **(D)** Superplots showing PCC between Itgb1 and Lamp1 in WT and Rab7KO mouse fibroblast. “P” indicated the p-value obtained by two-sided Welch’s *t*-test from the mean values of independent experiments, N=3. “p” indicated the p-value obtained by two-sided Welch’s *t*-test from all individual values collected, WT n=82, KO n=103 cells. Bars represent the Mean±SD of the mean values. **(E)** Quantification of EGFP intensity differences before and after NH_4_Cl treatment in individual WT and Rab7KO mouse fibroblast. Values were measured on cell bodies in processed images and normalized to the pretreatment condition. P-value was analyzed by two-sided Welch’s *t*-test. WT n=34, KO n=35 cells from 4 independent experiments. Bars represent Mean±SD. **(F-G)** Quantification of EGFP-positive (F) and EGFP/Sir-Lysosome (SirLyso) double positive (G) structures in WT and Rab7KO mouse fibroblasts following the NH_4_Cl treatment. Statistics was analyzed by two-sided Welch’s *t*-test. WT n=34, KO n=35 cells from 4 independent experiments. Bars represent Mean±SD. **(H)** Quantification of the ratio of SirLyso-positive versus EGFP/SirLyso double positive structures in WT and Rab7KO mouse fibroblasts following the NH4Cl treatment. P-value was analyzed by two-sided Welch’s *t*-test. Cells without double positive structures were omitted from the analysis. WT n=21, KO n=35 cells from 4 independent experiments. Bars represent Mean±SD.

**Figure S3.**
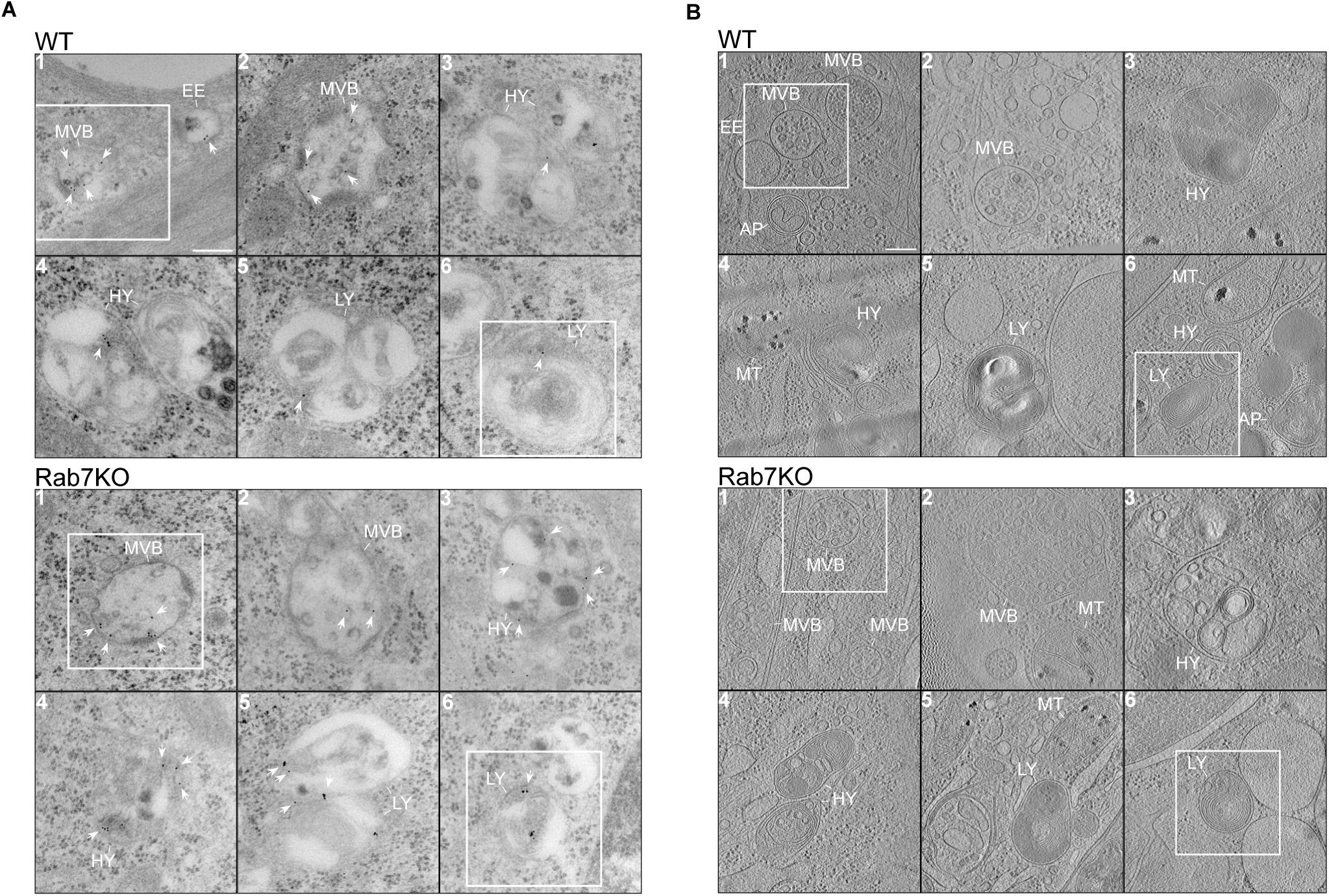
Related to Figure 2. **(A)** TEM images of Itgb1 containing endosomes and lysosomes in the WT and Rab7KO mouse fibroblast. EE, early endosome, MVB, multi-vesicular body; HY, hybrid endosome (endolysosome or autolysosome); LY, lysosome. Arrowheads indicate immunogold-labelled Itgb1. Scale bar, 0.2 µm. n=2 independent experiments with at least 2 different EM grids analyzed. Boxes indicate area shown in Figure 2E. **(B)** Cryo-EM images of acidic endosomes and lysosomes in WT and Rab7KO mouse fibroblast. EE, early endosome, MVB, multi-vesicular body; AP, autophagosome; HY, hybrid endosome (endolysosome or autolysosome); LY, lysosome; MT, mitochondria. Scale bar, 0.2 µm. WT n=3, KO n=2 independent experiments with at least 2 different EM grids analyzed. Boxes indicate area shown in Figure 2F, corresponding tomographs are shown in Video S1.

**Figure S4.**
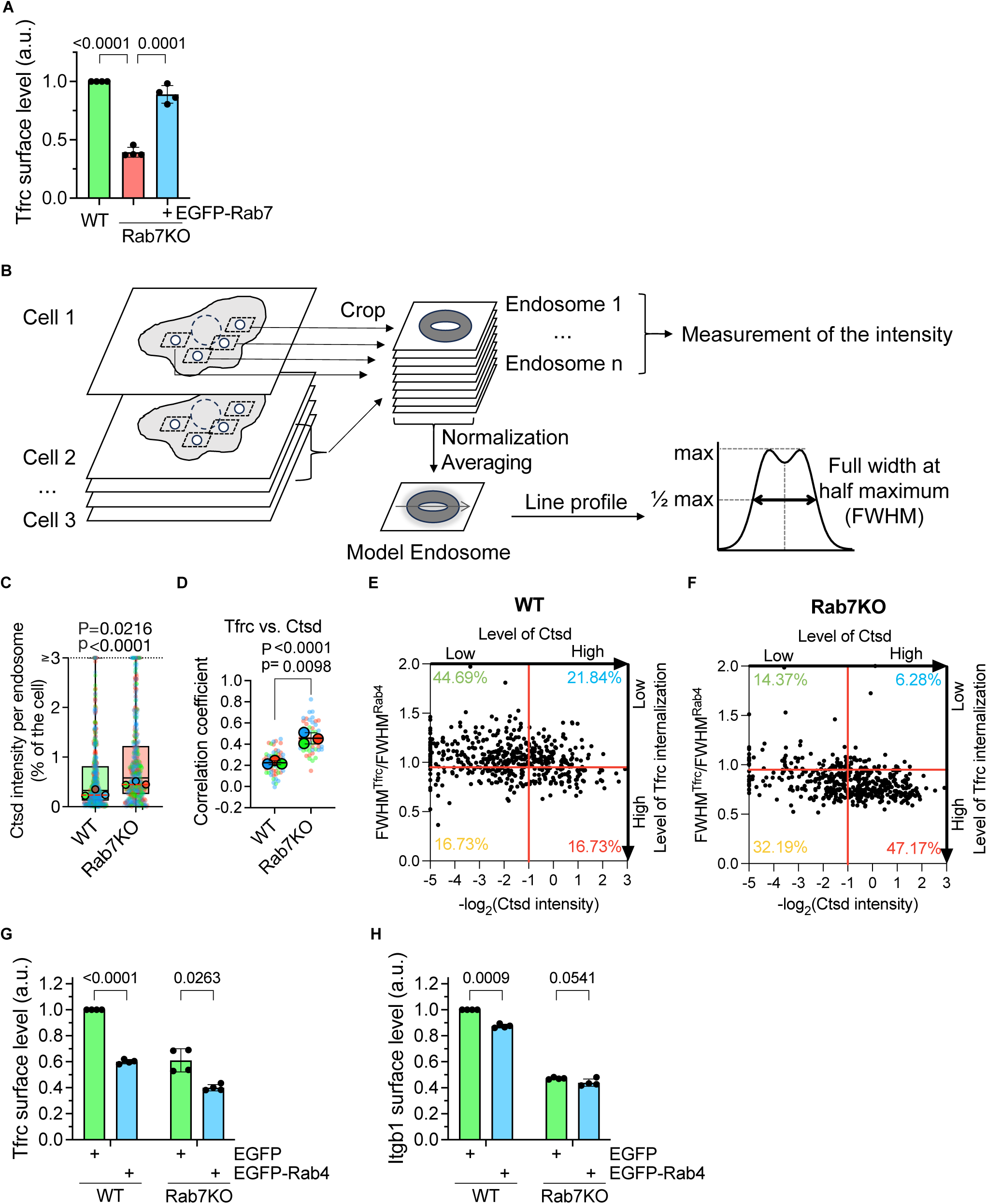
Related to Figure 4. **(A)** Cell surface levels of Tfrc in WT and Rab7KO fibroblasts, and Rab7KO fibroblasts stably re-expressing EGFP-Rab7 determined by flow cytometry. Statistics was analyzed by one sample *t*-test when compared to WT or by two-sided Welch’s *t*-test. Data are shown as Mean±SD, n=4 independent experiments. **(B)** Schematic of the workflow for quantitative analysis of recylolysosomes. Enlarged Rab4^+^ endosomes with a donut-like appearance were collected from sum-projected confocal stacks. To measure the relative accumulation of fluorescence signal on endosome, the pixel values were transformed from grey level to the percentage of intensity in each endosome compared to the whole cell. To assess the full width at half maximum (FWHM) of the fluorescent signals, endosomes were normalized for intensity and isotropy and averaged to obtain a model endosome, on which a line profile could be generated. **(C)** Quantification of the Ctsd intensity of individual endosomes. The integrated intensity was measured in a round area with a diameter of 1.4μm in the center of the image. On top of box-and whisker plot with all individual values shown, dots with solid outline show the geometric means of each independent experiment, red dashed lines show the mean value of 3 geometric means. Values higher than 3 were plotted on the border. “P” indicated the p-value obtained by two-sided Welch’s *t*-test from the geometric means of each independent experiment, N=3. “p” indicated the p-value obtained by Mann-Whitney test from all individual values collected, WT n=516, KO n=505 endosomes. **(D)** Superplots showing PCC between Tfrc and Ctsd in WT and Rab7KO mouse fibroblast expressing EGFP-Rab4. “P” indicated the p-value obtained by two-sided Welch’s *t*-test from the mean values of each independent experiment, N=3. “p” indicated the p-value obtained by two-sided Welch’s *t*-test from all individual values collected, WT n=62, KO n=58 cells. Bars represent the Mean±SD of the mean values. **(E-F)** Quantification of Tfrc internalization and Ctsd accumulation in individual enlarged Rab4^+^ endosomes of WT (E) and Rab7KO (F) fibroblasts expressing EGFP-Rab4. The y-axis indicates the ratio between FWHM of Tfrc signals and Rab4 signals in individual endosomes. The x-axis indicates the -log_2_ transformed Ctsd intensity of individual endosomes. The red lines indicate the cutoffs between the high and low groups, arbitrarily set at x=-1 and y=0.95. Numbers indicate the percentage of endosomes classified in each group. Endosomes with erroneous FWHM values (negative, extremely large) due to the presence of interfering signals from a second structure were omitted from the analysis. Points with x-values less than 5 were plotted along the border. Y-axis is capped at y=2. WT n=490, KO n=494 endosomes. **(G-H)** Cell surface levels of Tfrc (F) and Itgb1 (G) determined by flow cytometry in WT and Rab7KO fibroblasts transiently expressing EGFP or EGFP-Rab4. Statistics was analyzed by two-sided multiple paired *t*-test with Holm-Šidák correction. Data are shown as Mean±SD, n=4 independent experiments.

**Figure S5.**
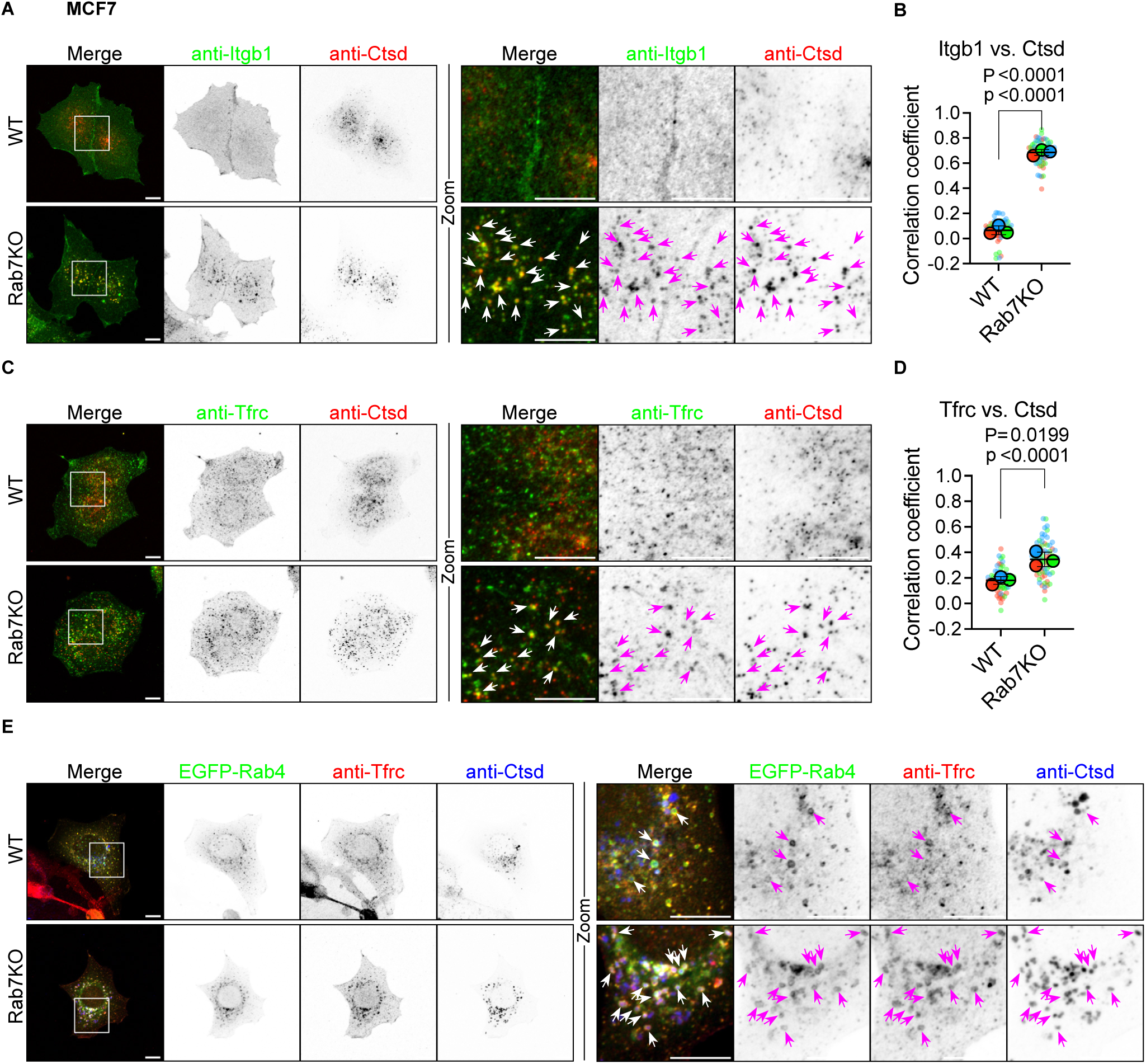
Related to Figure 4. **(A)** Representative IF images of Itgb1 and Ctsd in WT and Rab7KO MCF7 cells. Boxes indicate cytoplasmic areas shown in Zoom. Sum intensity projections of confocal stacks are shown. Arrowheads indicate Itgb1 accumulation in Ctsd-positive lysosomes. Scale bar, 10µm. **(B)** Superplots showing PCC between Itgb1 and Ctsd in WT and Rab7KO MCF7 cells. “P” indicated the p-value obtained by two-sided Welch’s *t*-test from the mean values of each independent experiment, N=3. “p” indicated the p-value obtained by two-sided Welch’s *t*-test from all individual values collected, WT n=67, KO n=71 cells. Bars represent the Mean±SD of the mean values. **(C)** Representative IF images of Tfrc and Ctsd in WT and Rab7KO MCF7 cells. Boxes indicate cytoplasmic areas shown in Zoom. Sum intensity projections of confocal stacks are shown. Arrowheads indicate Tfrc accumulation in Ctsd-positive lysosomes. Scale bar, 10µm. **(D)** Superplots showing PCC between Tfrc and Ctsd in WT and Rab7KO MCF7 cells. “P” indicated the p-value obtained by two-sided Welch’s *t*-test from the mean values of each independent experiment, N=3. “p” indicated the p-value obtained by two-sided Welch’s *t*-test from all individual values collected, WT n=67, KO n=74 cells. Bars represent the Mean±SD of the mean values. **(E)** Representative IF images of Tfrc and Ctsd in WT and Rab7KO MCF7cells expressing EGFP-Rab4. Boxes indicate cytoplasmic areas shown at in Zoom. Sum intensity projections of confocal stacks are shown. Arrowheads indicate colocalization of Rab4 and Ctsd. Scale bar, 10µm.

**Figure S6.**
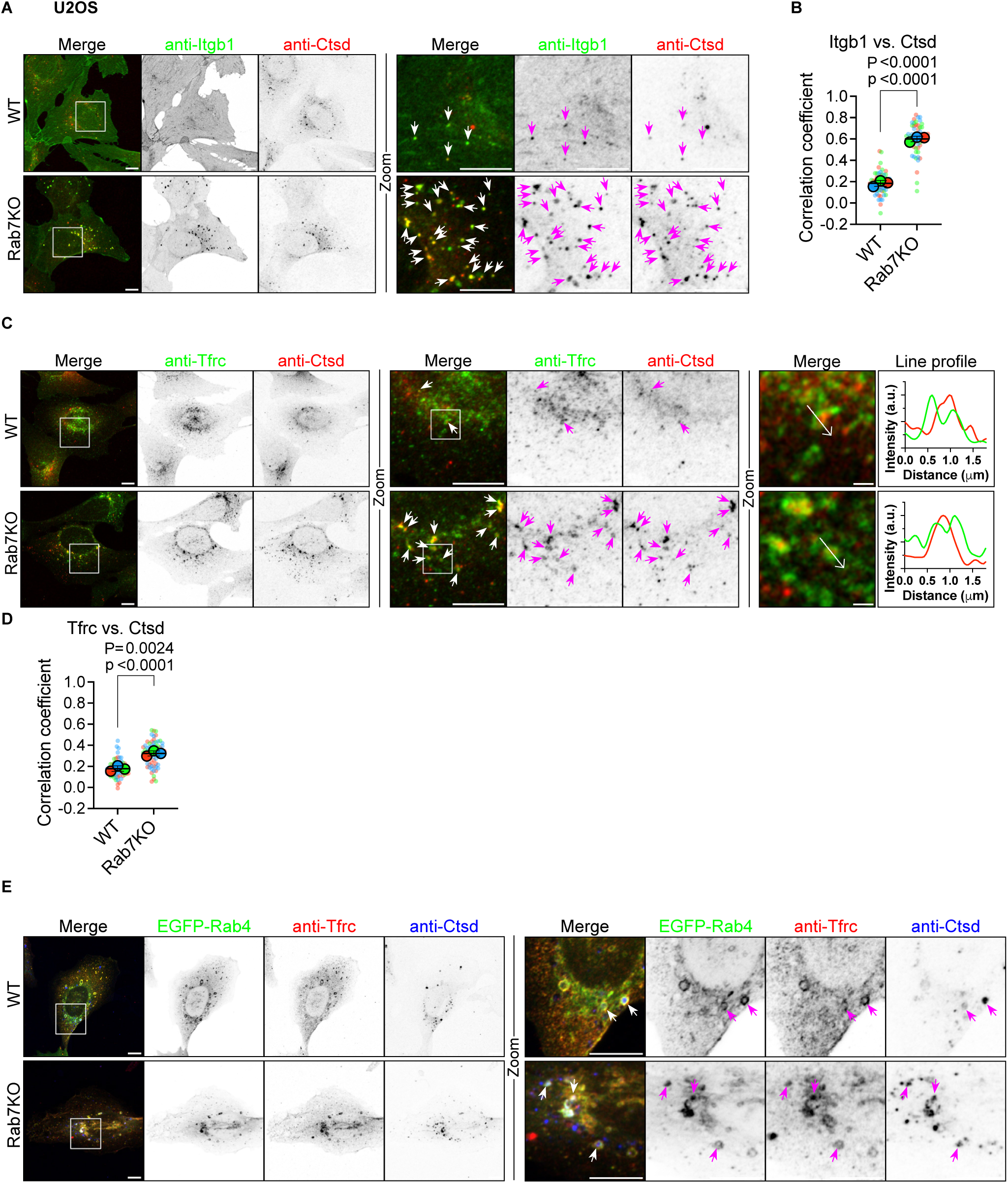
Related to Figure 4. **(A)** Representative IF images of Itgb1 and Ctsd in WT and Rab7KO U2OS cells. Boxes indicate cytoplasmic areas shown in Zoom. Sum intensity projections of confocal stacks are shown. Arrowheads indicate Itgb1 accumulation in Ctsd-positive lysosomes. Scale bar, 10µm. **(B)** Superplots showing PCC between Itgb1 and Ctsd in WT and Rab7KO U2OS cells. “P” indicated the p-value obtained by two-sided Welch’s *t*-test from the mean values of each independent experiment, N=3. “p” indicated the p-value obtained by two-sided Welch’s *t*-test from all individual values collected, WT n=65, KO n=57 cells. Bars represent the Mean±SD of the mean values. **(C)** Representative IF images of Tfrc and Ctsd in WT and Rab7KO U2OS cells. Boxes indicate cytoplasmic areas shown in Zoom. Sum intensity projections of confocal stacks are shown. Arrowheads indicate Tfrc accumulation in Ctsd-positive lysosomes. Arrows in single confocal slice indicate direction of line profiles of Tfrc (green) and Ctsd (red). Scale bar: left and middle panel 10µm, right panel 1μm. **(D)** Superplots showing PCC between Tfrc and Ctsd in WT and Rab7KO U2OS cells. “P” indicated the p-value obtained by two-sided Welch’s *t*-test from the mean values of each independent experiment, N=3. “p” indicated the p-value obtained by two-sided Welch’s *t*-test from all individual values collected, WT n=61, KO n=66 cells. Bars represent the Mean±SD of the mean values. **(E)** Representative IF images of Tfrc and Ctsd in WT and Rab7KO MCF7cells expressing EGFP-Rab4. Boxes indicate cytoplasmic areas shown in Zoom. Sum intensity projections of confocal stacks are shown. Arrowheads indicate colocalization of Rab4 and Ctsd. Scale bar, 10µm.

**Video S1**

Cryo-tomography of endosomal organelles in WT and Rab7KO mouse fibroblast. Scale bar, 0.2µm.

## Resources table

### Antibodies

2 Antibodies
| Name | Manufacture | Cat. No. | Dilution | RRID |
| --- | --- | --- | --- | --- |
| **Immunofluorescence** |  |  |  |  |
| mouse Itgb1 | Homemade<sup>1</sup> |  | 1:5000 |  |
| mouse Cathepsin B | R&Dsystems | AF965 | 1:200 | RRID:AB_2086949 |
| mouse Cathepsin D | R&Dsystems | AF1029 | 1:200 | RRID:AB_2087094 |
| mouse Cathepsin L | R&Dsystems | AF1515 | 1:200 | RRID:AB_2665930 |
| mouse LAMP1 | DSHB | 1D4B | 1:250 | RRID:AB_2134500 |
| mouse LAMP2 | DSHB | ABL-93 | 1:100 | RRID:AB_2134767 |
| Transferrin receptor | ThermoFisher | 13-6800 | 1:250 | RRID:AB_2533029 |
| human Cathepsin D | R&Dsystems | AF1014 | 1:200 | RRID:AB_2087218 |
| human Itgb1 (TS2/16) | Biolegend | 303002 | 1:500 | RRID:AB_314317 |
| **Western blot** |  |  |  |  |
| mouse Itgb1 | Homemade<sup>1</sup> |  | 1:10000 |  |
| human Itgb1 | BD Biosciences | 610467 | 1:1000 | RRID:AB_2128060 |
| Rab7 (D95F2) | Cell Signaling | 9367 | 1:1000 | RRID:AB_1904103 |
| Gapdh | Calbiochem | CB1001 | 1:10000 | RRID:AB_2107426 |
| beta Tubulin | DSHB | E7 | 1:2000 | RRID:AB_528499 |
| LC3 | Sigma | L8918 | 1:2000 | RRID:AB_1079382 |
| **ELISA** |  |  |  |  |
| Itgb1 (MB1.2) | Sigma | MAB1997 | 1:250 | RRID:AB_2128202 |
| **Flow cytometry** |  |  |  |  |
| human Itgb1 (TS2/16) PE | Biolegend | 303004 | 1: 400 | RRID:AB_314319 |
| mouse Itgb1 (HMβ1-1) PE | Biolegend | 102207 | 1: 400 | RRID:AB_312885 |
| mouse CD71 (RI7217) APC | Biolegend | 113820 | 1: 100 | RRID:AB_2728135 |
| IgG mouse isotype control PE | Biolegend | 400114 | 1: 400 |  |
| IgG hamster isotype control PE | Biolegend | 400907 | 1: 400 | RRID:AB_326593 |
| **Immunoprecipitation** |  |  |  |  |
| anti-GFP | Sigma | 11814460001 | 1:1000 | RRID:AB_390913 |
| **Secondary antibodies** |  |  |  |  |
| Donkey anti-mouse A488+ | ThermoFischer | A32766 | 1:400 | RRID:AB_2762823 |
| Donkey anti-rat A488 | ThermoFischer | A21208 | 1:400 | RRID:AB_2535794 |
| Donkey anti-goat A568 | ThermoFischer | A11057 | 1:400 | RRID:AB_2534104 |
| Anti-rabbit FluoTag-X4 AF647 | Nanotag | N2404-AF647-S | 1:250 |  |
| Donkey anti-rat A647+ | ThermoFischer | A48272 | 1:250 | RRID:AB_2893138 |
| Donkey anti-rabbit 6nm immunegold | Jackson | 711-195-152 | 1:40 | RRID:AB_2340609 |
| Goat anti-rabbit 5nm immunegold | Sigma | G7277 | 1:40 | RRID:AB_259951 |
| Rhodamine Phalloidin | ThermoFischer | R415 | 1:500 |  |

### Reagents

3 Reagents
| Name | Manufacture | Cat. No. |
| --- | --- | --- |
| Protamine sulphate | Sigma | P4020 |
| Hoechst 33343 | Thermofisher | H1399 |
| PfuUltra II DNA Polymerases | Agilent | 600674 |
| Dynabeads M-270 Streptavidin | Thermofisher | 65305 |
| Circligase II | Lucigen | CL9025K |
| Precision gRNA Synthesis Kit | Thermofisher | A29377 |
| TrueCut Cas9 Protein | Thermofisher | A36498 |
| DMSO | Sigma | D8418 |
| Bafilomycin A1 | Adipogen | BVT-0252 |
| MG132 | Sigma | 474787 |
| EZ-Link Sulfo-NHS-SS-Biotin | Thermofisher | A39258 |
| EZ-Link Sulfo-NHS-LC-Biotin | Thermofisher | 21335 |
| Streptavidin Mag Sepharose | Cytiva | 28985799 |
| iST - LYSE buffer | PreOmics | P.O.00032 |
| Exosome Immunoprecipitation Reagent (Protein G) | Thermofisher | 10612D |
| Prolong Glass | Thermofisher | P36984 |
| Fibronectin | MerckMillipore | 341631 |
| SiR-Lysosome | Spirochrome | SC012 |
| LysoTracker Deep Red | Thermofisher | L12492 |
| Dynabeads MyOne carboxylic acid | Thermofisher | 65011 |

## Methods

### Cell culture

Hap1 cells were obtained from Horizondiscovery (#C631, RRID:CVCL_Y019) and cultured in IMDM (Gibco#31980030) with 10% FBS (Gibco#A5256701) and SV40 Large T-immortalized mouse fibroblasts previously described^2^. MCF7 (#HTB-22, RRID:CVCL_0031) and U2OS (#HTB-96, RRID:CVCL_0042) cells were from ATCC. HEK237T cells were obtained from Takara Bio (#632273, RRID:CVCL_B0XW). All cell lines were cultured in DMEM (Gibco#61965059) with 10% FBS (Gibco#A5256701) at 37°C with 10% CO_2_ and regularly checked for mycoplasma.

### Expression constructs

pGT-GFP^3^ was a gift from Thijn R. Brummelkamp (Netherlands Cancer Institute, The Netherlands). To generate stable expression construct of the Cas9 protein, eSpCas9Plus (a gift from Ervin Welker, Addgene# 126767, RRID:Addgene_126767) was cloned into a pLIX-based lentiviral vector. To generate transient sgRNA expression constructs, a U6 promoter and an spCas9 sgRNA platform were cloned from PX459V2 (a gift from Feng Zhang, Addgene#62988, RRID:Addgene_62988) into a pMAX(Lonza)-based vector. The guide sequences were then cloned into the vector using the same cloning protocol as for PX459. To generate stable expression construct of human Rab7, the EGFP-Rab7A (a gift from Qing Zhong, Addgene#28047, RRID:Addgene_28047) was cloned into a pRetroQ (Clontech)-based retroviral vector. Retroviral expression construct of Human Itga5-EGFP has been described previously ^2^. pEGFP-Rab4A was a gift from Marci Scidmore (Addgene# 49434, RRID:Addgene_49434). pEGFP-Rab5A and pEGFP-Rab7A were gifts from gift from Guido Serini (Torino University, Italy). To generate stable expression construct of Rab4, mGreenLantern^4^ sequence and mouse Rab4A sequence (CCDS52703.1) were synthesized and cloned into a pT4 (Addgene#108352, RRID:Addgene_108352)-based Sleeping Beauty gene expression vector. EGFP-Rab5 sequence was cloned from pEGFP-Rab5 into the Sleeping Beauty vector. hsSB^5^ sequence was synthesized and cloned into a pMAX(Lonza)-based expression vector.

### Whole genome screen

The HAP1 haploid screen was performed as previously described with slight modifications^3^. Hap1 cells were seeded into 10 T175 flasks at 1.6 million cells/flask. The next day, cells were incubated with the gene-trapping retrovirus for 48 hours in the presence of 8ug/ml protamine sulfate. Mutagenized HAP1 cells were maintained in culture for two weeks, and 400 million cells were subcultured at each passage. 2.5 billion cells were harvested from the last culture and resuspended to 100 million cells/ml in FACS buffer (PBS containing 2% FBS and 2.5 mM EDTA). Cells were stained with PE-labeled anti-Itgb1antibody (Biolegend#303004) for 45 minutes on ice, washed twice with ice-cold PB, fixed with BD fixation buffer (BD#554655) for 10 minutes on ice and further incubated for 10 minutes at room temperature. Fixed cells were stored in FACS buffer supplemented with 0.01% sodium azide at 4°C in the dark before sorting on a FACSAriaIII flow cell sorter. Prior to loading onto the sorter, cells were stained with Hoechst 33343 (Thermofisher #H1399) for 30 minutes on ice. Cells were first gated on the Hoechst channel for 1n DNA content, then on the FSC and SSC channels for singlet, and finally on the PE channel to collect the 5% high and 5% low Itgb1 expressing cell populations.

Collected cells were lysed overnight at 56°C with agitation and genomic DNA was extracted using the NucleoSpin Blood L kit (Macherey Nagel# 740954). For each population, 8 linear PCRs were set up using Agilent PfuUltraII DNA polymerase (Agilent#600674), each containing 2ug of genomic DNA and 0.75 pmol of biotinylated primer (5’-[biotin-TEG]GGTCTCCAAATCTCGGTGGAAC-3’) in a volume of 50ul and running for 120 cycles. Every 2 PCR reactions were pooled and purified in 100ul elution buffer using Monarch PCR & DNA Cleanup Kit (7:1 binding buffer, NEB#T1030L). 5ul/reaction Dynabeads M270 Streptavidin (Thermofisher#65305) was washed in PBS+0.05% Triton and resuspended in 50ul/reaction 2× Binding Buffer (6 M LiCl, 10 mM Tris, 1 mM EDTA, pH 8.0, 0.1% Triton). The PCR elution was added to the binding buffer containing the beads to a final volume of 200ul. ssDNA was pooled for two hours at room temperature. ssDNA bound to the beads was washed in PBS containing 0.05% Triton. An adapter sequence for Illumina sequencing was ligated to the 3’ of the ssDNA on the beads using Cricligase II (Lucigen#CL9025K). A mixture containing 12.5 pmol ssDNA linker (5’-[phospho]CTGTCTCTTATACACATCTCCGAGCCCACGAGACACTCA[dideoxycytidine]-3’) 2.5 mM MnCl_2_, 1 M betaine, 1 µl 10X reaction buffer and 0.5 μl of Circligase II in a total volume of 10 µl was used for each linear PCR reaction. Ligation was performed at 60°C for 2 hours. Beads were washed three times with PBS+0.05% Triton and used as template for PCR to add Illumina i7 indexed barcodes using primers 5′-AATGATACGGCGACCACCGAGATCTACAC<u>ATCTGATGGTTCTCTAGCTTGCC</u>-3′ and 5’-CAAGCAGAAGACGGCATACGAGAT [i7 index] <u>GTCTCGTGGGCTCGG</u>-3′. Products from four PCR reactions were pooled and purified using NucleoSpin Gel and PCR Clean-up Kit (Macherey-Nagel #740609). Equal amounts of products were mixed and sequenced on Illumina NextSeq 500 instruments using the NextSeq high-output kit (75 cycles) and the sequencing primer 5’-CTAGCTTGCCAAACCTACAGGTGGGGTCTTTCA-3’.

The fastq output files were analyzed on the Galaxy platform^6^ using the instances at usegalaxy.org and usegalaxy.eu. Reads were mapped to the human genome hg38 using BWA-MEM. Sense insertions into each gene were counted using featureCounts with a customized Ensembl GRCh38 GTF file that includes only the non-UTR regions of protein-coding genes.

For each gene, the mutation index (MI) was calculated using the formula below:

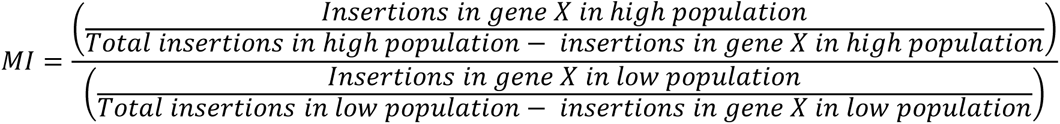

Entities without a valid HGNC symbol or without any insertions were omitted. Genes without a single insertion in either population were assigned a value of 1 so as not to be omitted from the plots. P-values were determined using the Chi-squared test and corrected for false discovery rate using the BH method. Data were analyzed and visualized using R with the packages tidyverse, ggplot and ggrepel.

### Generation of knockout cell lines

To generate Rab7A KO mouse fibroblasts, sgRNAs were generated with the targeting sequence CGACAGACTTGTTACCATGC. To generate Rab7A KO HAP1 cells, sgRNAs were generated with the targeting sequences AATCAGTACAAAGCCACAAT. Cells were reverse transfected with 3ug TrueCut Cas9 (Thermofisher#A36498), 1ug pEGFP-C1 plasmid, 0.6 µg *in vitro* transcribed sgRNA (Precision gRNA Synthesis Kit, Thermofisher#A29377) using Lipofectamine CRISPRMAX (Thermofisher#CMAX00003). GFP+ cells were sorted at 1 cell/well into 96 well plates 24 hours after transection. Western blot and sequencing were used to confirm the KO clones. To generate Rab7KO HEK293T, MCF7 and U2OS cell lines, cells were first transduced with lentivirus to express the eSpCas9Plus-T2A-TagBFP2 protein. One plasmid encoding a GFP protein and two plasmids expressing sgRNA (targeting sequence AATCAGTACAAAGCCACAAT and TGACAGGCTAGTCACAATGC) were transiently transfected into the cells. To generate a superclone, 100 GFP+ & BFP+ cells were sorted into one well of a 96-well plate two days after transfection. Western blot was used to confirm the KO. Unless explicitly indicated otherwise, experiments were conducted using clone or superclone number 2.

### Transfection

To generate stable pGT-GFP, EGFP-Rab7A, Itga5-EGFP and eSpCas9Plus expressing cell lines, cells were transduced overnight in 6-well plates with viral particles concentrated from cell culture supernatant as previously described^7^. For transient expression of EGFP-Rab4A, EGFP-Rab5A or EGFP-Rab11A expression plasmids, cells were reverse transfected with 1ug plasmids in a 6-well plate using JetOptimus (Polyplus#101000025) or Lipofectamine2000 (Thermofisher#11668019) following the manufacturer’s protocol. To generate cell lines stably expressing GFP-tagged Rab4A and -Rab5A, cells were reverse transfected with 0.5ug pT4-mGreenLantern-Rab4A or pT4-EGFP-Rab5A and 0.1ug hsSB plasmids in a 6-well plate using JetOptimus.

### Western blot

Cells were lysed in lysis buffer containing 50 mM Tris-HCl, pH8.0, 150 mM NaCl, 5 mM EDTA, 1% NP-40 and cOmplete tablet (Roche#4693159001). For drug treatment, cells were incubated with bafilomycin A1 (10 nM) or MG132 (20 µM) overnight prior to the lysis. BCA assays were used to determine protein concentrations. 5 ug protein samples were boiled at 90°C for 5 minutes in LDS sample buffer (Genscript#M00676 or Millipore#MPSB) containing 5% 2-mercaptoethanol, processed for SDS-PAGE (Genscript SurePAGE precast gel or Millipore mPage precast gel) and transferred to PVDF membranes (Millipore#IPVH00010). After blocking with 3% BSA or 5% milk in TBST, the membranes were blotted with primary antibodies and then HRP-conjugated secondary antibodies. Chemiluminescence detection (Immobilon Western Chemiluminescent HRP substrate, Millipore#WBKLS0500) was performed using a GE Amersham AI600 imager. All experiments were repeated at least three times with similar results.

### Quantification of surface Itgb1 level

Cells were detached from the culture plates using Accutase (Sigma#A6964), washed in ice-cold PBS and stained with antibodies diluted in 100ul FACS buffer (PBS containing 2% FBS and 2.5 mM EDTA) for 45 minutes on ice. For drug treatment, cells were incubated with bafilomycin A1 (10 nM) or MG132 (20 µM) overnight before detachment. After two ice-cold PBS washes, surface levels of Itgb1 were quantified using an LSRFortessa X-20 Cell Analyzer. FlowJo software was used for data analysis. Experiments were repeated at least three times with similar results.

### Cell proliferation assay

Mouse fibroblasts were seeded onto glass bottom ibidi µ-Dish 35 mm (ibidi#81158) coated with 10 µg/ml FN. Cells were washed twice with DMEM 6 hours after adherence and the imaging dish was mounted on a Nikon Ti2 microscope. Phase contract images were captured at 1-hour intervals for at least 24 hours using a Plan Achromat 10X objective. During imaging, cells were maintained in DMEM with 10% FBS and 20 mM HEPES at 37°C and with 5% CO2. Cells were manually counted at 0h and 24h in the same fields of view using FIJI’s multi-point tool. The difference in cell number was then used to calculate cell doubling time. Data were obtained from three independent experiments.

### Cell adhesion assay

The cell adhesion assay was performed as previously described^8^. In brief, cells were grown to 70-80% confluence, serum starved for at least 4 hours and detached by trypsin/EDTA. 30,000 cells were seeded into 96-well plates coated with 3% BSA or 0.01% poly-L-lysine (PLL) or 5 µg/ml FN in PBS for the indicated time points and washed twice by immersion in PBS before fixation with 4% PFA. Adherent cells were then stained with crystal violet (0.1% in 20% methanol) and washed with tap water. Stained cells were solubilized with 2% SDS and read on a plate reader at a wavelength of 595 nm and the results were normalized using the following formula: Normalized OD_595_= (OD_FN_-OD_BSA_)/(OD_FN_-OD_PLL_). Data were obtained from three independent experiments.

### Cell spreading assay

The cell spreading assay was performed as previously described^8^. In brief, 50,000 serum-starved cells were allowed to spread on 5 ug/ml FN-coated 15 mm coverslips for the indicated time points. Cells were washed with PBS and fixed with 4% PFA before staining with rhodamine phalloidin (ThermoFischer #R415) and Hoechst 33342 (ThermoFischer #H1399). Cells were then imaged using a Leica LSM780 confocal microscope and the spreading area of at least 50 cells per condition per experiment was quantified using ImageJ. Data were obtained from four independent experiments.

### Itgb1 Cycloheximide chase assay

2.5×10^5 cells per well were cultured overnight in a 6-well plate prior to cycloheximide treatment (at a final concentration of 5 ug/ml). At the indicated time points, cells were lysed in lysis buffer (50 mM Tris, 150 mM NaCl, 0.1% SDS, 0.1% SDC, 1% Triton, 5mM EDTA and cOmplete tablet) and total protein was quantified by BCA assay. The same amount of protein from each sample was loaded for WB to determine Itgb1 and Gapdh levels. Data were obtained from four independent experiments.

### Itgb1 surface stability assay

Turnover of surface Itgb1 were measured as previously described^2^. In brief, 2.5×10^5 cells per well were cultured overnight in a 6-well plate and surface labeled with NHS-LC biotin (Thermofisher#21335) at a concentration of 0.2 mg/ml in PBS for 20 minutes on ice. Cells were then incubated at 37°C for the indicated time points before lysis in ELISA lysis buffer (50 mM Tris, 150 mM NaCl, 0.5% NP-40, 1.5% Triton, 5mM EDTA and cOmplete tablet). Cell lysate was added overnight to an ELISA plate precoated with anti-Itgb1 antibody (Sigma#MAB1997), the plates were then washed and the Itgb1 levels measured upon addition of HRP-streptavidin using a microplate reader at 405 nm with ABTS substrate. Data were obtained from four independent experiments.

### Itgb1 internalization assays

The internalization rate of Itgb1 was measured as previously described^9^. In brief, cells were surface-labeled with NHS-SS-Biotin (Thermo Scientific# A39258) and incubated over different time points before MesNa (Sodium 2-mercaptoethanesulfonate, Sigma# M1511) reduction. Cells were then lysed in ELISA lysis buffer (50 mM Tris, 150 mM NaCl, 0.5% NP-40, 1.5% Triton, 5mM EDTA and cOmplete tablet) and a capture ELISA was performed. The amount of Itgb1 measured corresponds to the amount internalized in the cell. Data were obtained from four independent experiments.

### Cell surface proteome

Mouse fibroblasts were grown to 70% confluence in three independent 6 cm dishes. Cells were placed on ice, washed twice with ice-cold DPBS, and incubated in surface biotinylation buffer (0.2 mg/ml sulfo-NHS-SS-biotin in DPBS, Thermofisher#A39258) for 45 minutes at 4°C. Cells were washed twice with ice-cold PBS and lysed in lysis buffer containing 50 mM Tris-HCl, pH 8.0, 150 mM NaCl, 5 mM EDTA, 1% NP-40 and cOmplete Tablet. Streptavidin Mag Sepharose beads (Cytiva#28985799) were used to pull down biotinylated proteins for 1 hour at 4°C. Beads were washed twice with lysis buffer, then twice with PBS to remove detergent. On-beads bound proteins were incubated for 20 mins at 37°C with 30 µL of 1x SDC buffer (1% sodium deoxycholate (SDC), 40 mM 2-Cloroacetamide (CAA), 10 mM TCEP in 100 mM Tris, pH 8.0). After 1:1 dilution with MS grade water, the proteins were digested overnight at 37°C by addition of 0.5 µg of trypsin. With the help of a magnetic rack, the supernatant was separated from the beads and collected. After acidification with Trifluoroacetic acid (TFA) to a final concentration of 1%, peptides were desalted using in-house produced strong cation exchange (SCX) stage tips.

Desalted peptides were applied onto a 30-cm column (inner diameter: 75 microns; packed in-house with ReproSil-Pur C18-AQ 1.9-micron beads, Dr. Maisch GmbH) at 60°C using the autosampler of the Thermo Easy-nLC 1200 (ThermoFisher). Eluted peptides were directly sprayed onto the timsTOF Pro (Bruker Daltonics). Peptides were loaded at 400 nl/min in buffer A (0.1% FA) and percentage of buffer B (80% acetonitrile, 0.1% FA), ramped from 5% to 25% over 90 minutes followed by a ramp to 35% over 30 minutes, a ramp to 58% over 5 minutes, final to 95% over the next 5 minutes and maintained at 95% for another 5 minutes.

TimsControl was used to perform data acquisition on timsTOF Pro. The mass spectrometer was set to data-dependent PASEF mode and performed one survey TIMS-MS and ten PASEF MS/MS scans per acquisition cycle. Analysis was performed in a mass scan range from 100-1700 m/z and an ion mobility range from 1/K0 = 0.85 Vs cm-2 to 1.35 Vs cm-2 using equal ion accumulation and and dual TIMS analyzer ramp time of 100 ms each at 9.43 Hz spectra rate. Suitable precursor ions for MS/MS analysis were isolated in a window of 2 Th for m/z < 700 and 3 Th for m/z > 700 by rapid switching of the quadrupole position in sync with the elution of precursors from the TIMS instrument. Collision energy was decreased as a function of ion mobility from 45eV for 1/K0 = 1.3 Vs cm-2 to 27eV for 0.85 Vs cm-2. Collision energies were linearly interpolated between these two 1/K0 values and maintained constant above and below these base points. A polygon filter mask was utilised for eliminating singly charged precursor ions, and additional m/z and ion mobility information was employed for ‘dynamic exclusion’ to avoid re-sequencing of precursors that attained a ‘target value’ of 14500 a.u.

### Total cell secretome

Mouse fibroblast cells were grown to 70% confluence in DMEM with serum in three independent 15 cm dishes. Cells were washed twice with DPBS and then incubated in 10 ml DMEM+20 mM HEPES without serum for 7.5 hours at 37°C. The culture medium was collected and centrifuged at 2000g for 20 minutes and then filtered through a 0.45um syringe filter. Proteins were precipitated by adding trichloroacetic acid (TCA) to a final concentration of 10%, incubated overnight at 4°C, and pelleted at 5200 g for 1 hour at 4°C. The supernatant was removed and the pellets were washed twice with 5 mL of cold 100% ethanol and air dried for 20 minutes at room temperature. Resuspensions were made by adding 100 µl of PreOmics LYSE buffer (PreOmics#P.O.00032) to the pellets, boiling for 5 minutes at 90°C with agitation, and sonicating for 10 minutes at room temperature in a water bath sonicator. Samples were diluted at a 1:1 ratio with MS grade water and digested overnight at 37 °C with 1 µg LysC and with 2 µg trypsin. The peptide solution was then acidified with TFA to a final concentration of 1%, followed by desalting with SCX stage tips.

Desalted peptides were loaded onto a column as described above. Eluted peptides were directly sprayed onto the QExactive HF mass spectrometer (ThermoFisher). Peptides were loaded at 400 nl/min in buffer A (0.1% FA) and percentage of buffer B (80% acetonitrile, 0.1% FA), ramped from 7% to 30% over 60 minutes followed by a ramp to 60% over 15 minutes, 95% over the next 5 minutes and maintained at 95% for another 5 minutes. The MS was operated in a data-dependent mode with survey scans from 300 to 1650 m/z (resolution of 60000 at m/z =200), and up to 10 of the top precursors were selected and fragmented using higher energy collisional dissociation (HCD) with a normalized collision energy value 28. MS2 spectra were acquired at a resolution of 15k at m/z = 200. AGC target for MS1 and MS2 scans were set to 3E6 and 1E5 respectively within a maximum injection time of 100 and 60 ms for MS and MS2 scans respectively.

### Endosome-IP and proteomic profiling

Mouse fibroblast cells expressing GFP-tagged Rab4A or Rab5A were grown to 70% confluence in three independent 15 cm dishes. Cells were rinsed three times in ice-cold DPBS before scraping from the plate into 1 ml homogenization buffer (HB) containing 250 mM sucrose, 20 mM HEPES, 10 mM imidazole, 5 mM EDTA, and 0.03 mM cycloheximide. Cells were centrifuged at 400g for 5 minutes at 4°C. The supernatant was removed, and the pellets were gently resuspended in 1 ml HBC buffer (HB with cOmplete tablet) using a wide-cut pipette tip. The suspension was loaded into a 1 ml syringe and passed through a G26 needle 10 to 20 times. The process was monitored by phase contrast microscopy by adding 1ul of homogenate to 15ul of 0.04% trypan blue in PBS. Homogenization was stopped when the plasma membranes of 70%-80% of the cells ruptured. Homogenates were centrifuged at 2000g for 10 minutes at 4°C. The supernatants were collected and centrifuged again at 2000g for 10 minutes at 4°C. The supernatants after the second centrifugation were collected and referred to as postnuclear supernatant (PNS). For each IP sample, 5 µg anti-GFP antibody (Roche#11814460001) was coupled to 50 µl Protein G Dynabeads slurry (Thermofisher#10612D) according to the manufacturer’s instructions. Beads were washed twice with PBS to remove detergent before being added to PNS to pull down intact endosomes at 4°C for 1 hour. Beads were washed three times with HB and boiled in 100 µl PreOmics LYSE buffer (PreOmics#P.O.00032) at 90°C for 10 minutes. The supernatants were collected, diluted 1:1 with water, and digested overnight at 37°C by addition of 0.5 µg of LysC and 1 µg of trypsin. After acidification with TFA (final concentration of 1%), peptides were desalted using SCX stage tips.

Desalted peptides were loaded onto a column as described above. Peptides were separated by an Easy-nLC 1200 (Thermo Fisher Scientific) coupled to an Exploris 480 mass spectrometer (Thermo Fisher Scientific) at a flow rate of 300 nL/min. The analytical column was heated to 60 °C. As gradient, the following steps were programmed with increasing addition of buffer B (80% Acetonitrile, 0.1% formic acid): linear increase from 5 to 30% over 40 minutes, followed by a linear increase to 95% over 10 minutes and finally, the percentage of buffer B was maintained at 95% for another 10 minutes. The mass spectrometer was operated in data-dependent mode with survey scans ranging from m/z 300 to 1650 Th (resolution of 60k at m/z = 200 Th), and up to 15 of the most abundant precursors were selected and fragmented using stepped Higher-energy C-trap Dissociation (HCD with a normalized collision energy of value of 30). MS2 spectra were acquired using dynamic m/z range (resolution of 15k at m/z = 200 Th). The normalized AGC targets for the MS1 and MS2 scans were set to 300% and 100%, respectively. The maximum injection time for the MS1 scan was 25 ms. For MS2 scans, the maximum injection time was set to auto.

### MS data analysis

Raw data were processed using the MaxQuant computational platform^10^ (version 1.6.7.0 or 2.0.1.0) with Orbitrap or TimsTOF data using standard settings. The peak list was searched against the Uniprot database of Mus Musculus (55466 entries, July 2020). Cysteine carbamidomethylation was set as a static modification, and methionine oxidation, deamidation and N-terminal acetylation were set as variable modifications. The match-between-run option was enabled, and proteins were quantified across samples using the label-free quantification algorithm in MaxQuant, which generates label-free quantification (LFQ) intensities.

Perseus software^10^ (v.1.6.15.0) was used to perform statistical analysis on the LFQ data. LFQ intensities were log_2_ transformed. Proteins that were known contaminants, reverse, only identified by site or had less than 3 valid values were removed. Missing values were imputed using normal distribution (width 0.3, down shift 1.8). The significance of differences in the cell surface proteome and cell secretome between cell lines was estimated using the Perseus’ Volcano plot function, and the adjusted p-value was calculated using a two-sided permuted *t*-test (250 randomizations, FDR=0.05 and S0=0.1).

To compare differential protein enrichment on endosomes, a volcano plot comparing Rab4 and Rab5 groups for each cell lines were first done, and adjusted p-values were obtained using two-side permuted *t*-test (250 randomizations, FDR=0.05). Then the overall difference between the WT and Rab7KO cells lines for each protein were calculated using the following formula:

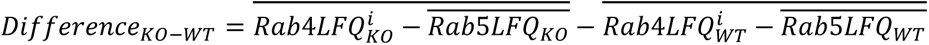

Where *LFQ*^*i*^was the LFQ value of protein in each independent experiment and 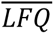 was the mean value three independent experiments. The significance of differences was estimated using the Perseus’ Volcano plot function, and the adjusted p-value was calculated using a two-sided permuted *t*-test (250 randomizations, FDR=0.05 and S0=0.1). Visualization of the data was done using R scripts with the Tidyverse and ggrepel packages.

### Gene ontology analysis

Gene ontology enrichment analysis was conducted utilizing the clusterProfiler^11^ R package. P-value was adjusted using the BH method. Figures were generated using the ggplot2 package.

### Immunofluorescence

Cells were grown overnight on glass coverslips coated with 10 µg/ml FN. Cells were fixed with 4% PFA for 15 minutes at room temperature, permeabilized with 0.1% Triton for 4 minutes, and blocked with blocking buffer (1x ROTI block in PBS, CarlRoth#A151) for 1 hour at room temperature. Cells were incubated with primary antibodies diluted in blocking buffer overnight at 4°C and with secondary antibodies for 1 hour at room temperature. Coverslips were mounted in Prolong Glass (ThermoFisher# P36984), dried at room temperature for 5 days, and stored at 4°C until imaging. Z-stacks were acquired using a Zeiss LSM 980 confocal microscope with Airyscan2 and a Zeiss Plan-Apochromat 63X/1.40 NA oil immersion objective. Colocalization analysis was performed using the EzColocalization^12^ plug-in in FIJI. Prior to processing, images were subjected to Gaussian filter-based background subtraction as described below (radius=0.61um). Pearson’s correlation coefficient was calculated above the Costes threshold. Experiments were repeated at least three times with similar results.

### NH_4_Cl neutralization assay

Mouse fibroblast expressing Itga5-EGFP were grown overnight on ibidi µ-Slide 8 Well high Glass Bottom coated with 10ug/ml FN. Cells were incubated with SirLysosome for 1 hour and washed twice with and kept in 200 µl live cell imaging solution (140 mM NaCl, 2.5 mM KCl, 1.8 mM CaCl_2_, 1.0 mM MgCl_2_, 20 mM HEPES, 4.5 g/ml glucose, and pH=7.4). Time-lapsed z-stacks were recorded at 1 minute and 1 µm intervals by using a Nikon Eclipse Ti2 microscope, equipped with a Princeton Instruments ProEM EMCCD camera and an Okolab microscopy cage incubator, with a Plan Apochromat 60X/1.40 NA oil-immersion objective. 200 µl of 2X NH_4_Cl buffer (100 mM NH_4_Cl, 40 mM NaCl, 2. 5mM KCl, 1.8 mM CaCl2, 1.0 mM MgCl2, 20 mM HEPES, 4.5 g/ml glucose) were injected into the imaging chamber during time-lapse recording. Images were denoised and subjected to a Gaussian-filter-based background subtraction (radius=0.65um) as described below. Z-stacks were projected to 2D using the “Z Projection-Max Intensity” function in FIJI. Experiments were repeated at least three times with similar results. To quantify the number of EGFP-positive and Sir-Lysosome positive structures, the cell body area was enclosed in an ROI and the default “enhance contrast” command (resulting saturation = 0.35) was applied to each channel. Fluorescent puncta were manually quantified using the point tool in Fiji.

### Image denoising

A CARE(2D) machine learning-based denoising model^13^ was trained using ZeroCostDL4Mic^14^. To generate training data, images of fixed cells stained with anti-Itgb1 antibody and Alexa Fluor 488 secondary antibody were acquired on the above-mentioned live cell microscope. Low signal/noise (S/N) images were acquired with an exposure time of 10 ms, while high S/N images were acquired at the same stage position with an exposure time of 50 ms. The trained model was applied to the images in FIJI using the CSBDeep plugin.

### Gaussian-filter-based background subtraction

A background image was created from the input image using the Gaussian Blur function in FIJI. The background was then subtracted from the input image using the ImageCalculator function.

### Endosome maturation flux assay

Enlarged Rab4^+^ endosomes with a visually donut-like appearance in the EGFP channel and with no adjacent endosome present were cropped from the SUM-projected confocal stacks. The geometric center of the donut was manually adjusted. For each endosome, the original image was duplicated 359 times and each replica was rotated one degree with respect to the previous one. All 360 images were stacked and z-projected to obtain an isotropic image. The value of pixels was normalized to the range of 0-1 based on the maximum and minimum value within a circular area of 2 µm in diameter. To generate a model endosome, all endosomes were then combined into a stack and average-projected. Line profiles were made at the center line of the picture. A 6th degree polynomial interpolation was used to provide an accurate estimation of the full width at half maximum (FWHM). The FWHM of individual endosomes was determined in the same way. To measure Ctsd filling, the intensity of each pixel was set as a percentage of the integrated intensity of the entire cell. The integrated intensity was then measured in a circular ROI of 1.4 µm in diameter. Cells were collected from three independent experiment.

### Transmission Electron Microscopy

Mouse fibroblasts were seeded and grown overnight on ACLAR® Film 0.2 mm thickness (Ted pella#10501) coated with 10 µg/ml FN in a 24-well plate. Cells were rinsed twice with ice-cold DMEM containing 20 mM HEPES and incubated for 30 minutes on ice with anti-Itgb1 antibody diluted in DMEM containing 10% FBS and 20mM HEPES. After three washes with ice-cold DMEM-HEPES, cells were incubated for 30 minutes on ice with 6 nm (Jackson#711-195-152) or 5 nm (Sigma#G7277) diameter immunogold coupled secondary antibodies in DMEM-FBS-HEPES. After three washes in ice-cold DMEM-HEPES, cells were incubated in DMEM-FBS-HEPES at 37°C for 2 hours to allow internalization of the immunogold labeled integrin. Specimens were fixed with 2% glutaraldehyde (Electron Microscopy Sciences #16220) in DMEM-HEPES for 45 minutes at room temperature and then overnight at 4°C.

Samples were washed two times with 0.1 M sodium cacodylate (Electron Microscopy Sciences#11653) pH 7.4 and post-fixed with 1% OsO_4_ (Electron Microscopy Sciences#19193) in 0.1 M sodium cacodylate pH 7.4, for 40 min. After two washes with distilled water, samples were gradually dehydrated by successive baths in 30%, 50%, 70%, 90%, 96%, and 100% ethanol. Samples were embedded in Spurr’s Low Viscosity Embedding Media Kit (Electron Microscopy Sciences#14300) following the manufacturer’s protocol. Small columns of polymerized resin were placed on the Aclar-foil with a drop of resin between cells and column. Samples were polymerized at 60° for 48 hours. A pyramid (100×500 µm) was trimmed with Leica Ultramicrotome EM UC6 and trimming knife. Ultra-thin serial sections (60 nm) were cut with Ultramicrotome EM UC6 and stained with Leica Ultrostainer with 0.5% Uranyl acetate (Electron Microscopy Sciences#22400) and Ultrostain-2 3% Lead citrate (Leica Microsystems#16707235). Images acquisition was done with JEOL JEM-1230 transmission electron microscope, 80kV, with Gatan Orius SC1000 digital Camera and Gatan Digital Micrograph™ software. Data were collected from two independent experiments and each time at least two different grids.

### Correlative light-electron tomography

Cells were seeded and grown overnight on Quantifoil carbon film gold grids (Quantifoil R 1/4 on 200 gold mesh) coated with 10 µg/ml FN. Cells were rinsed twice with DMEM and labelled for 5 minutes at 37°C with Lysotracker Deep Red diluted in DMEM. Grids were taken out by plunge tweezer and extra medium on grid was removed by filter paper. 4 µl cryo-buffer contain DMEM, Lysotracker, 5% glycerol and 50x Dynabeads MyOne (ThermoFisher#65601) was applied on the grid, then blotted and plunged in ethane–propane with a Vitrobot Mark IV (blot time 10s, blot force 8, room temperature, 100% humidity). Grids were stored in liquid nitrogen until further image acquisition.

Grids were autoclicked with custom cut-out autogrids, and loaded onto a Leica SP8 cryo-confocal microscope equipped with a cryo-stage and a 50x 0.9NA objective (Leica Objective No. 506520). Stacks were acquired using a 638-nm laser for Lysotracker and fiducial beads autofluorescence, along with brightfield images. After deconvolution using Huygens Essential (https://svi.nl/Huygens-Software), the stacks were resliced to isotropic pixel size using the 3D-correlation toolbox (3DCT) (https://3dct.semper.space/). To prepare the lamellae, the grids were then transferred to the ThermoFisher Cryo-FIB/SEM Aquilos microscope. A protective CpMePtMe3 layer was applied on the grid using Gas injection system. Then SEM and ion beam images were acquired for each target square. 3D correlation of SEM images with fluorescence images was performed based on the fiducial beads in 3DCT. The Lysotracker+ areas in 3DCT were marked for the lamella milling. At the target position, milling was performed simultaneously from above and below step by step: 0.3 nA until 1.2 µm, 0.1 nA until 0.8 µm, for rough milling, and 50 pA until 250 nm, 30 pA until 100 nm for fine milling. To confirm that target organelles were still remained in the lamella, grids were reloaded on cryo-fluorescent microscopy to check Lysotracker signals when the lamella thickness reached 0.8 µm. Positions displayed Lysotracker signals were selected to fine milling.

Grids were loaded into the TEM with the previous milling direction perpendicular to the tilt axis. Tomogram acquisition positions were determined by correlation of fluorescence data with TEM images of the grid squares containing lamellae (3DCT), followed by inspection of low magnification lamella images.Tilts series were acquired with Titan Krios (field emission gun 300 kV, Thermo Fisher Scientific) equipped with an energy filter and a direct detection camera (K2 Summit, Gatan) at a magnification of 42,000x (pixel size 3.52 Å) and defocus −5 µm. Frames were acquired in dose-fractionation mode with a total dose of 120 e−/A2 per tilt series using SerialEM 3.9.0 (https://bio3d.colorado.edu/SerialEM/). A dose-symmetric tilt scheme was used with an 2° increment in a total range of ±60° from a starting angle of 10° (+ or −) to compensate for lamella pretilt. Frames were aligned using MotionCorr2 (https://emcore.ucsf.edu/ucsf-software), and tomogram reconstruction was performed in IMOD (https://bio3d.colorado.edu/imod/). After 2x binning, tomograms reconstructed from odd/even frames were denoised using cryo-CARE^15^.

### Statistics

GraphPad Prism software was used for statistical analysis and graph tracing unless otherwise specified.

